# An optimized workflow for spatial transcriptomics across early development in Xenopus

**DOI:** 10.64898/2026.05.07.723548

**Authors:** Chenxi Zhou, Shubhamay Das, Thomas Defard, Kyra J. E. Borgman, Subham Seal, Vincent Kappès, Thomas Walter, Iva Simeonova, Geneviève Almouzni, Anne H. Monsoro-Burq

**Author notes:** equal contribution (Shubhamay Das, Thomas Defard).

## Abstract

How gene expression patterns change spatially as the embryo transitions from simple to complex structures remains a major developmental biology question. Recently developed imaging-based spatial transcriptomics (ST) enable mapping expression of multiple gene at a single-cell resolution. Although Xenopus is a key model in embryology there is no established ST pipeline, and commercially available techniques face many challenges (sample preparation, probe design, cell segmentation). Furthermore, the highly diverse cell shapes and sizes across developmental stages and between different tissues represent major hurdles to accurately defining cells. Here, we describe an optimized workflow for ST in blastula-to-tailbud-stage frog embryos using Merscope, commercial MERFISH (Multiplexed Error-Robust Fluorescence In Situ Hybridization) originally designed for standard mammalian tissues. With stringent quality control and tailored computational pipelines, we optimize this technology for robust, semi-quantitative profiling of spatial transcriptomic landscapes in non-mammalian embryos. Reliable tissue preservation and cell-segmentation enable high-resolution mapping of gene expression during the development of a complex multi-tissue organization. This versatile strategy applies broadly to various dynamic systems, from embryos of various model organisms to complex and heterogeneous organs in mammals.

**Summary statement:** This Single-cell Spatial Transcriptomics pipeline and reference atlas in *Xenopus -* a model organism in embryology - overcome technical challenges and resolve dynamic changes in patterning during development.

## Introduction

Rapid technical advances in the field of spatial transcriptomics (ST) have allowed mapping gene expression profiles within cells in their microenvironment. Applied to developmental biology, this could considerably advance our understanding of the finely coordinated developmental gene programs that pattern the early embryo and orchestrate morphogenesis (Chen et al., 2022).

ST technologies can be broadly classified into sequencing-based and imaging-based approaches (Rademacher et al., 2025). Sequencing-based ST, such as Visium (10x Genomics) and Stereo-seq (BGI), give access to the full transcriptome, but have limited spatial resolution (Chen et al., 2022; Oliveira et al., 2025). Imaging-based ST such as Merscope (Vizgen), Xenium (10x Genomics), and Molecular Cartography (Resolve Biosciences), provide high sub-cellular resolution down to 0.3 μm and detection of a selected panel of transcripts (Cadinu et al., 2024; Chen et al., 2015; Groiss et al., 2021; Janssens et al., 2025; Marco Salas et al., 2025; Rademacher et al., 2025). Consequently, the aim of the project orients the choice of the technology. For example, Merscope, which is based on multiplexed error-robust fluorescence in situ hybridization (MERFISH), aims to precisely measure the copy number of single RNA molecules without amplification bias with a sub-cellular resolution in a single-cell (Chen et al., 2015; Moffitt et al., 2018; Wang et al., 2018). MERFISH has many applications in adult tissues, such as visualizing brain organization or tumor complexity (Hara et al., 2021; Moffitt et al., 2018; Zhang et al., 2023). Imaging-based ST has thus become the gold standard for accessing, at a single-cell resolution, the dynamics of co-expression of selected genes of interest during spatial and temporal tissue patterning or to analyze tumor microenvironment complexity (Pozniak et al., 2025). However, commercially available gene panels and detection of transcripts with spatial information have only rarely been optimized for non-mammalian model organisms (Janssens et al., 2025; Wan et al., 2024). Smaller tissues such as early-stage embryos measuring only 1-2mm require specific adaptations for sample preparation and data analysis, as the most imaging-based techniques were initially conceived for large tissue slices from patients’ tumors or adult mouse tissues. Further, as cells greatly vary in size and shape with developmental stage and between distinct cell types and tissue layers, segmentation of individual embryonic cells within their tissue context face both general challenges and case specific difficulties.

A well characterized reference model in embryology for over 80 years is a tetrapod vertebrate, the amphibian Xenopus. Its external fertilization and development provide a unique access to the first hours and days of a whole organism and represents a system of choice for spatial transcriptomics applications, with developmental stage- and cell type-specific challenges. To capture the *in vivo* dynamics of developmental genes expression in early embryogenesis within an unperturbed entire organism, we applied Merscope/MERFISH to *Xenopus* frog embryos at multiple developmental stages, from the simply-organized blastula to the complex multi-tissue tailbud stage. Here, we set up a robust pipeline, from sample preparation to cell segmentation and gene clusters analysis (**Fig. 1**). We provide a protocol for preparation of high-quality thin formalin-fixed paraffin-embedded (FFPE) sections from *X. laevis* embryos, guidelines for mounting up to 50 embryos into the same histological block for simultaneous analyses in one experiment, and settings to integrate variation in embryonic shape (**Fig.2**). We also provide a strategy with two in-house Cellpose2 models trained on *X. laevis* tissues for optimized cell segmentation. Importantly, these consider histological variables, such as the presence of a blastocoel in early-stage embryos or differences in cell shape and size throughout development (**Figs 3-4**). Finally, we provide a spatially resolved expression reference atlas with a list of 54 developmental marker genes, to identify the main cell types at key embryo time points, including germ layer formation in gastrula and neural patterning at tailbud stage (**Fig. 5**).

**Figure 1:**
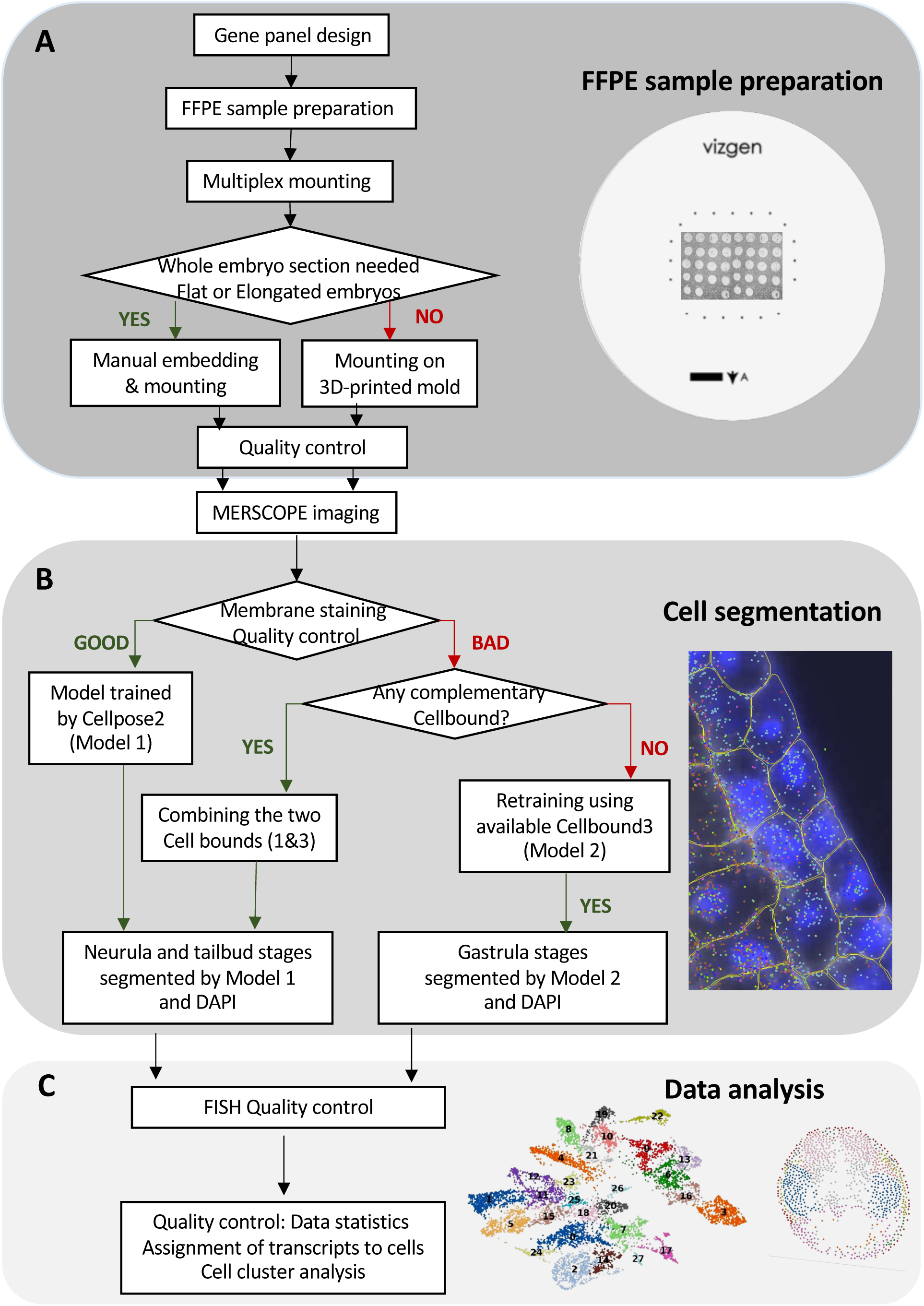
A workflow for Spatial Transcriptomics adapted to developing frog embryos. A. A custom gene panel was designed. Histology for early frog embryos was optimized. Up to 50 embryo sections were mounted onto a single imaging slide using two strategies depending on the shape (round or elongated) of the embryos. B. Following MERFISH hybridization and imaging, cell segmentation used a custom Cellpose2 model trained on *Xenopus laevis* sections, based on membrane and nucleus staining. According to specificities for each developmental stage and to the staining quality, the segmentation strategy was adapted (Model 1, Model2). C. Processed data were subjected to quality control and used in downstream analyses of gene expression and cell clustering.

**Figure 2:**
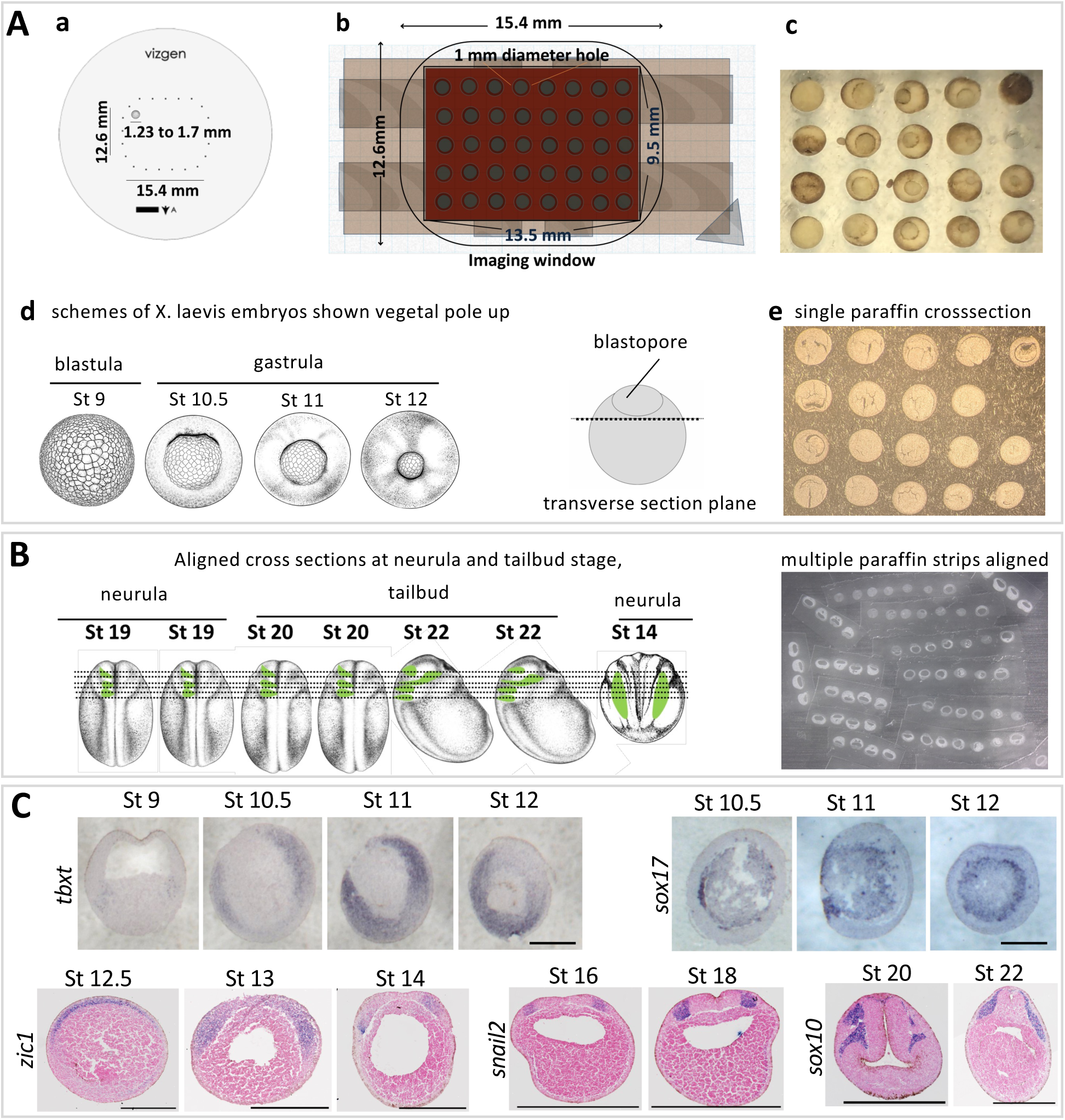
Optimization of *Xenopus laevis* embryo multiplexed sample preparation. A. (a). The relative size of the Vizgen Merscope slide v1 imaging window (dots) and of a *Xenopus laevis* embryo (grey circle) are compared. (b) Scheme of the 3D-printed mold designed to orient multiple round-shaped embryos and fit a single section on the slide. (c) Embryos mounted on top of the 3D-printed mold blastopore up or sideways according to need, during embedding in 70°C liquid paraffin. (d) Schemes of blastula- and gastrula-stage embryos and transverse section plane. (e) Two resulting paraffin cross sections of 20 embryos were fit on the v1 slide. B. Schematics of the manual alignment of several elongated embryos: the example shows seven neurula- (st 14, st 19) to tailbud- (st 20, st 22) stage embryos embedded together, dorsal up or sideways, aligning brain and cranial neural crest (in green). Transverse section planes are indicated. After close trimming of the paraffin, the resulting strips were mounted together on the Vizgen Merscope slide v1, here to obtain a total of 64 sections. C. After sectioning, RNA integrity was validated by colorimetric *in situ* hybridization on adjacent sections. Consistently high histology quality was obtained at all stages. *tbxt* labels the mesoderm and *sox17* the endoderm in gastrula stages 10.5, 11 and 12. At blastula stage 9 section, *tbxt* is barely detected in the marginal zone. In late gastrula (stage 12.5) and early neurula (stage 13, 14), *zic1* labels the neural ectoderm. At neurula stage 16, 18; *snail2* label neural crest cells. At tailbud stages 20, 22, *sox10* labels migrating neural crest cells. Optional eosin stain was used as a counterstaining of the cytoplasm. Scale bar: 500 μm.

**Figure 3:**
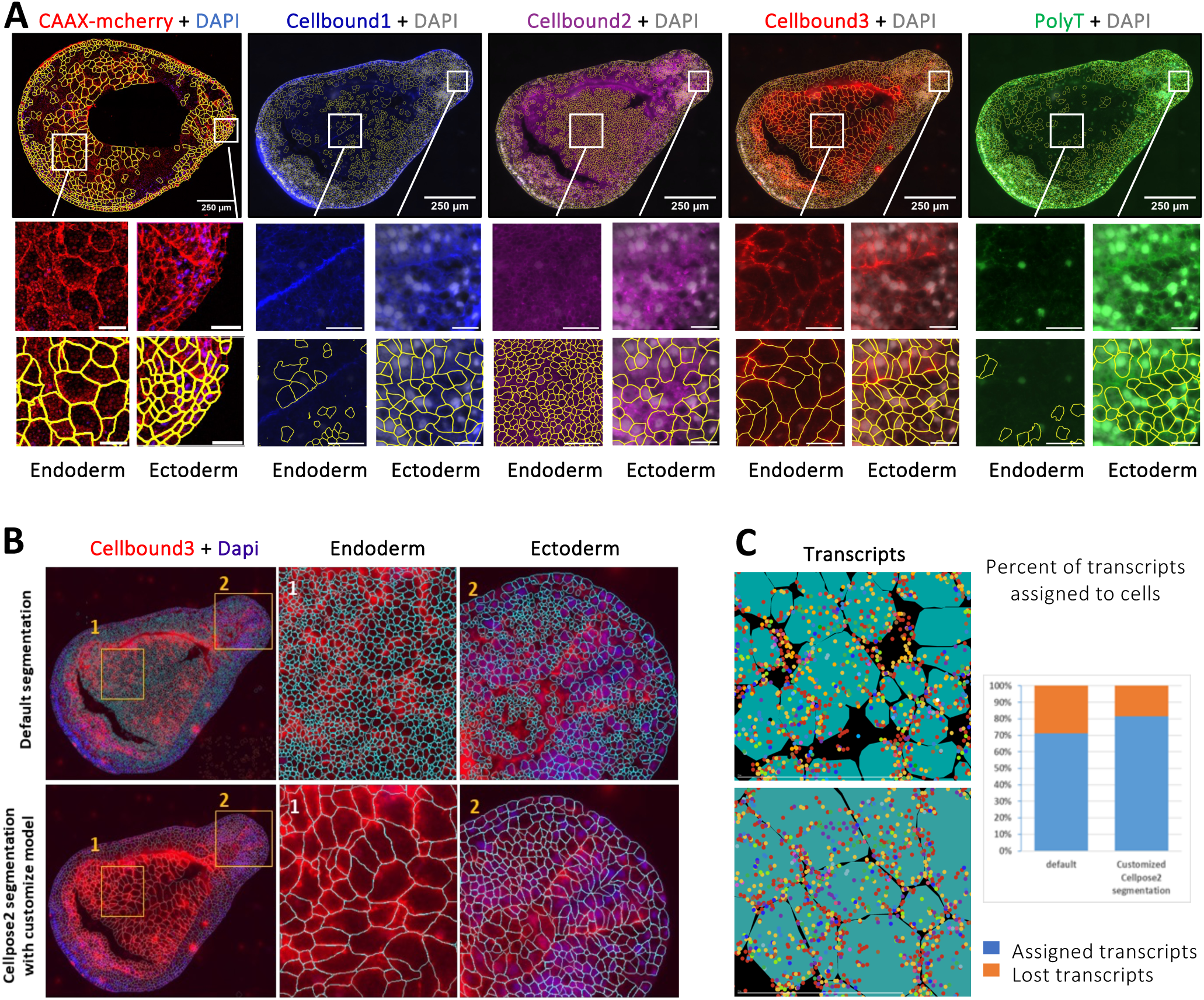
Cell segmentation using custom-trained Cellpose2, using Cellbound3 and DAPI staining, applied to tailbud stage 22 (Model 1). A. Cell segmentation (yellow lines) using the custom-trained Cellpose2 Model 1 with CAAX-mCherry, or based on membrane stains Cellbound1, Cellbound2, Cellbound3, PolyT, together with DAPI for nuclear stain. Cellbond3 + DAPI produced the cell segmentation most similar to CAAX-mcherry in both ectoderm and endoderm cells. Scale bars: upper panel/250μm, lower ectoderm/endoderm panels/25μm. B. Comparison of the default cell segmentation output (turquoise lines) and of Cellpose2 segmentation with the custom-trained Model 1. C. Custom cell segmentation retrieved a higher proportion of transcripts, compared to the default segmentation. Cells are as represented as turquoise surfaces and transcripts as colored dots.

**Figure 4:**
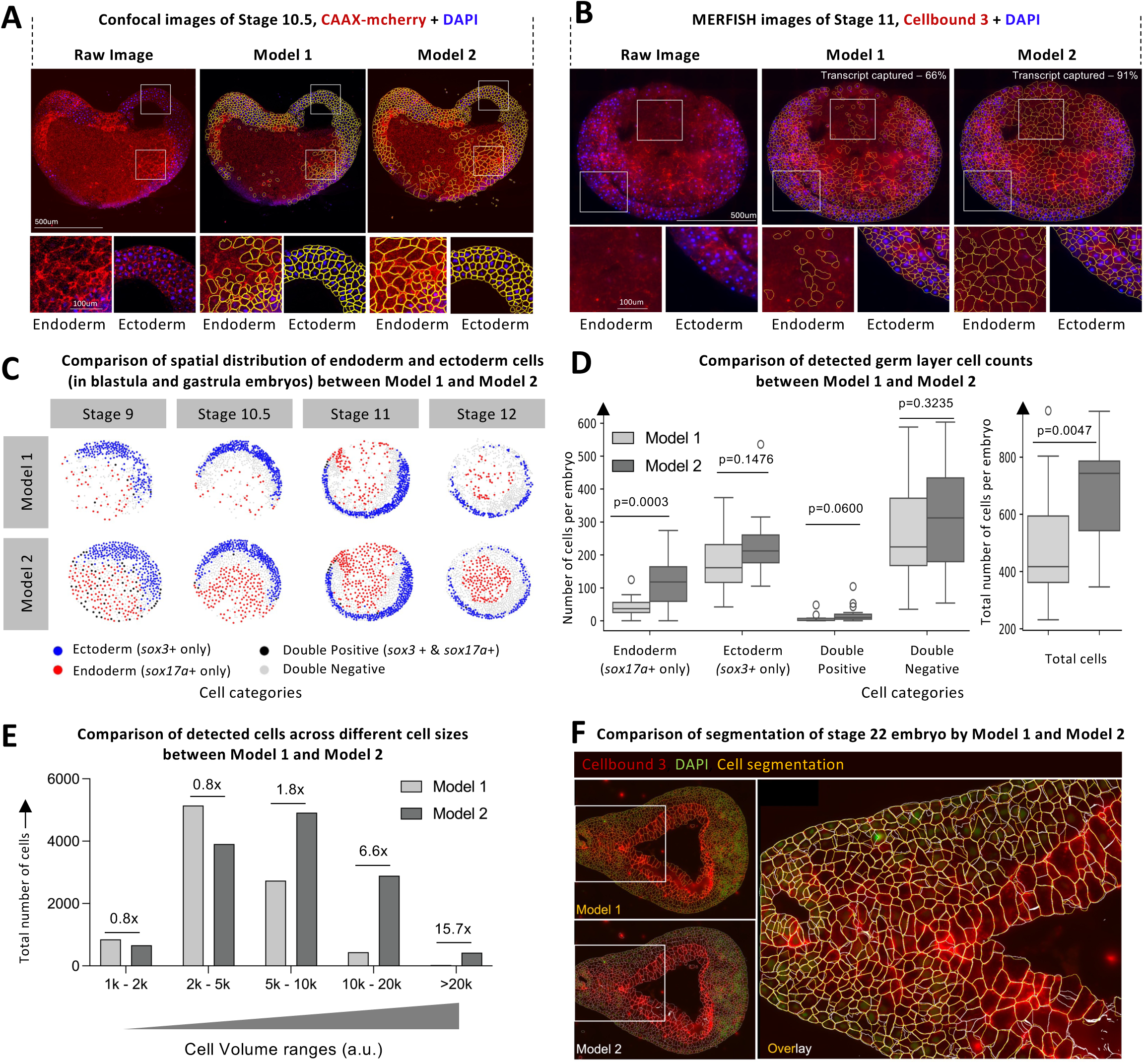
Early-stage embryos require additional segmentation optimization to identify specific cell types. A. Cell segmentation (yellow outlines) generated by stage-specific custom-trained Cellpose2 models using CAAX–mcherry and DAPI channels from confocal images of a stage 10.5 embryo. Model 1 was trained on Cellbound3 + DAPI images from stage 12.5 to 22 embryos as shown in figure 3, whereas Model 2 was trained on earlier stages (blastula stage 9 and gastrula stages 10.5, 11, and 12). Both models segmented ectodermal regions efficiently; however, in the endoderm region, Model 2 performed notably better by identifying a greater number of cells with more accurate and larger cell boundaries, while Model 1 tended to underestimate cell size and density. Scale bar of the upper panel: 500 μm, lower panel: 100 μm. B. Cell segmentation (yellow outlines) by Model 1 and Model 2, applied to stage 11 MERFISH images using Cellbound3 and DAPI channels. Model 2 achieved improved segmentation, particularly within the endodermal region. Scale bar of the upper panel: 500 μm, lower panel: 100 μm. C. Spatial distribution of endoderm and ectoderm cells detected by Model 1 and Model 2 in embryo sections from stages 9, 10.5, 11, and 12. Cells with ≥9 *sox3* transcripts were classified as ectoderm (blue dots), cells with ≥5 *sox17a* transcripts as endoderm (red dots), cells meeting both criteria as mixed identity (black dots), and cells meeting neither criterion as unclassified (grey dots). D. Quantification of endodermal and ectodermal cells detected per embryo by Model 1 and Model 2 in embryo sections from stages 9, 10.5, 11, and 12. Model 2 detected approximately three times as many *sox17a*-positive (endoderm) cells relative to the total cell count compared with Model 1, while detecting similar proportions of *sox3*-positive (ectoderm) cells. E. Quantification of cells detected across different cell sizes between Model 1 and Model 2. Model2 performed better in larger cell volumes and similarly in small to medium sized cells. F. Cell segmentation of a stage 22 embryo using Model 1 (yellow outlines) and Model 2 (white outlines) using Cellbound3 and DAPI channels. The strong overlap between yellow and white masks demonstrates that Model 2 retains high segmentation accuracy even at later developmental stages.

**Figure 5.**
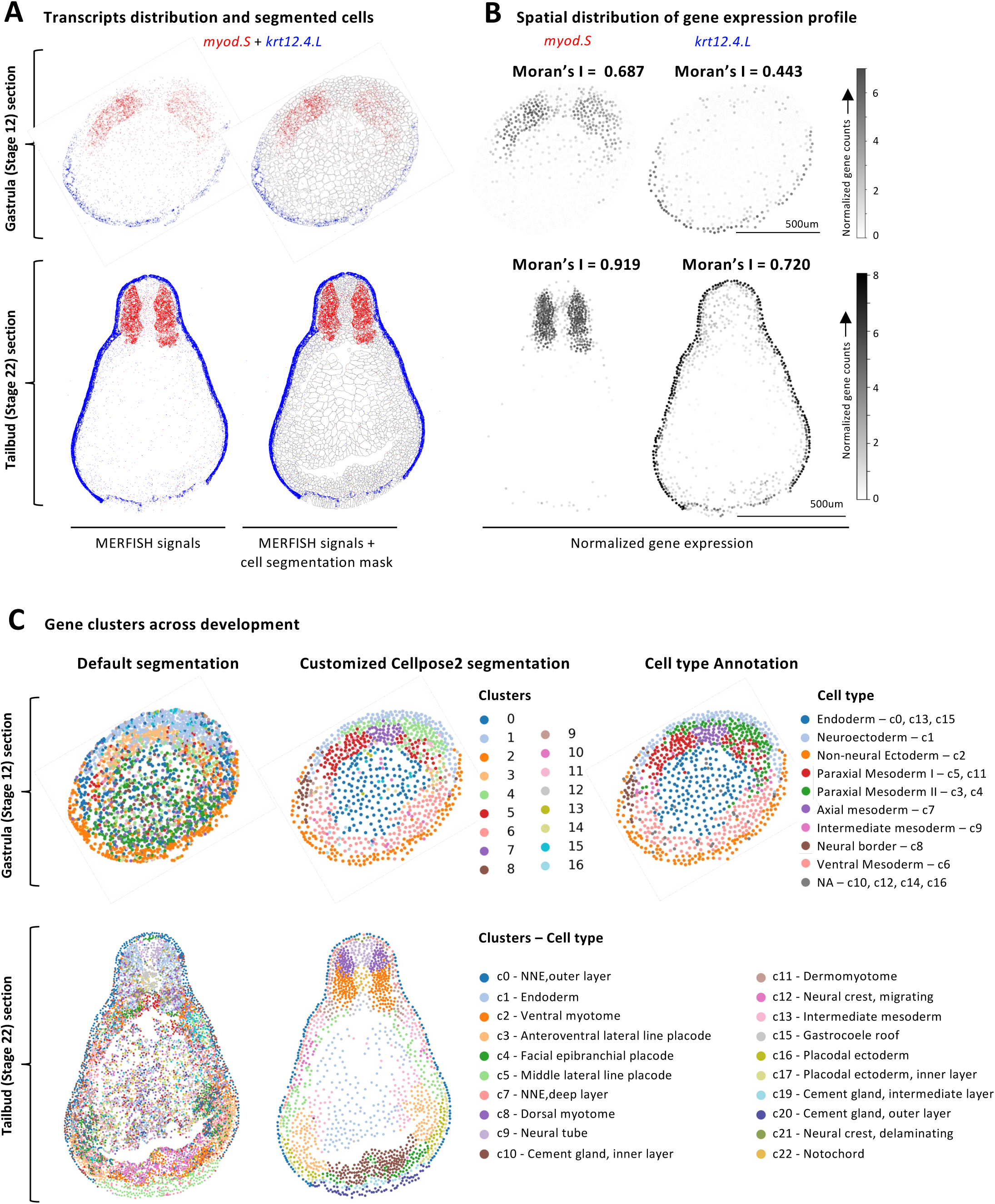
Transcript distribution and cell clusters at gastrulation and tailbud stages. A. Spatial transcript distribution of *myod.s* (red) and *krt12.4.L* (blue) on embryo sections at gastrula stage 12 and tailbud stage 22. Cell segmentation boundaries are in grey. B. Spatial expression pattern of *myod.s* and *krt12.4.L* in the same sections. Expression levels were calculated by volume-normalising and log-transforming transcript counts per cell, then represented by greyscale values within a cell. C. Spatial distribution of gene-based cell clusters in representative sections. Default Vizgen segmentation shows poor clustering. Customised Cellpose2 segmentation better segregated cells into spatially-defined clusters both at gastrula stage 12 (by Model 2) and tailbud stage 22 (by Model 1). At gastrula stage 12, we identified 8 main cell types – endoderm, neural ectoderm, neural border, non-neural ectoderm, axial mesoderm, intermediate mesoderm, paraxial mesoderm, ventral mesoderm, subdivided into 16 clusters according to their gene co-expression patterns (see text). At tailbud stage 22, we identified 10 main tissue types in the representative cranial sections – non-neural ectoderm, neural tube, neural crest, placodes, cement gland, notochord, myotome, dermomyotome, intermediate mesoderm, and endoderm, subdivided into 24 cell clusters, 22 being present on the example shown here. Cranial placodes comprised the *otx2+/rax+/sox2+/pax6+* optic placode (c6), the *otx2+/pax6+/cdh1+* lens placode (c25) and the *sox9+/sox10+/ephA2+* otic placode (c23), seen on other sections than the one depicted here. On this section are found the *sox3+/ephA4+/gbx2+* anteroventral lateral line placode (c3), the *six1+/eya1+/snai1+* facial epibranchial placode (c4), the *foxc2+/snai1+/eya1+/six1+* middle lateral line placode (c5), and two layers of early placodal tissue *ephA2+/snai1+/foxc1+* (c16) and *tfap2a+/cdh1+/eya1+* (c17).

Overall, our work provides a reliable pipeline for conducting ST analysis at early embryonic stages in a reference model organism. Our approach can apply to other vertebrate systems and potentially to *ex-vivo* models, such as “pseudoembryos” or tissue-specific organoids, to investigate gene expression and cell regulatory networks in normal or pathological development (Steventon et al., 2021).

## Results

### Formalin-fixed paraffin-embedded sample preparation for medium throughput mounting

Embryos that develop externally, including those of frogs, are yolk-rich and present large cavities such as the blastocoel at blastula and gastrula stages, or the archenteron at neurula and tailbud stages (**Fig. 2A-B**). To preserve tissue architecture in FFPE embryo samples, we found that the duration of each methanol dehydration and paraffin embedding step was crucial. Despite the small size of the embryos, a one-hour methanol wash as indicated in most current protocols (Wehrle et al., 2024) was insufficient, while an overnight final methanol wash was too long: both protocols rendered the embryos too brittle. In contrast, an overnight 70% methanol wash, followed by 3 hours of 100% methanol dehydration, reproducibly ensured high-quality histology in 7 μm-thick microtome sections, in both *X. laevis* and *X. tropicalis* embryos, across blastula to tailbud stages (**Fig. 2C, Fig. S1**). Although even with this improved protocol, we could not obtain consistently intact 5 μm-thick sections as in the commercial MERSCOPE protocol, 7 μm-thick sections gave satisfactory results. RNA integrity was assessed by colorimetric *in situ* hybridization and a Bioanalyzer DV200 above 80% (**Fig. 2C**, Monsoro-Burq, 2007; Materials and Methods; step-by-step protocol in Supplementary Materials).

Placing and orienting up to 50 embryo sections on a single spatial transcriptomics slide, initially designed for a single larger tissue section, required the implementation of two different strategies depending on the shape of samples – round or elongated. The imaging window of Vizgen MERSCOPE v1 slide measures 15.4mm×12.6mm, and 22mm×18mm for the Ultra v2 slide. Considering the average diameter of 1.2 to 1.7mm for a *X. laevis* embryo or tailbud stage (**Fig. 2A, b**, Leibovich et al., 2020), this provides the possibility to analyze multiple conditions together on the same slide, such as multiple developmental stages, or control and experimental conditions, minimizing batch effects and reducing costs. To do so, we increased paraffin temperature to 70 ℃ during the final embedding, to maintain paraffin liquid state throughout the time needed to precisely orient 40 to 50 embryos under a binocular microscope. This method preserved both histology and RNA quality (**Fig. 2C, Fig. S1).** Our first strategy was designed to embed multiple round-shaped embryos (blastula, gastrula and early neurula stage), by using a 3D-printed polymer mold with a grid of 5×8 1-mm diameter holes separated by 0.5 mm gaps. This maintained embryos in place and allowed paraffin access to embryos from all sides (**Figs 2A, S10**). One section of this block fit the v1 MERSCOPE slide (**Fig. 2A, e**). A limitation of this 3D-mold is that only the upper half of the embryo is accessible for sectionning. Embryos with various sizes or irregular/elongated shapes (such as late neurula or tailbud stage embryos) were difficult to align using the mold. The second strategy was then manual mounting and orientation in the 70℃-hot paraffin (**Fig. 2B**, Supplementary Materials). In both strategies, we prepared closely trimmed 7μm FFPE sections, that we mounted together as closely as possible, within the imaging window of the slide using traditional histology techniques (**Fig. 2B, 2A.e,** Supplementary Materials). The slide was then dried and processed according to the manufacturer’s instructions for nuclear and membrane staining, clearing and photo-bleaching to remove autofluorescence. It was then embedded in a gel, sequentially hybridized with the probes and imaged with the Merscope instrument (Vizgen).

### Cell segmentation with a custom Cellpose2 model using Cellbound3 and DAPI captures embryonic cell size and shape heterogeneity from mid-gastrulation stage to tailbud stages

The default Merscope output includes images of the nucleus (DAPI), total mRNAs (polyT) and cell membrane (Cellbound 1, 2, 3) staining, the coordinates of detected transcripts, the cell to gene matrix, and the cell metadata (including the cell volume and coordinates; **Fig. 3**). To produce the cell to gene matrix, the sections should first be segmented accurately into individual cells and transcripts assigned to each cell. However, the default cell segmentation was unsatisfactory our conditions, due to the large variation in cell size in a given embryonic section and to the inability of the algorithm to distinguish cells from yolk-platelets (**Fig. 3B**). Moreover, the cell membrane staining was critical for accurate annotation of endoderm cells, as due to their large volume, nuclear staining was missed in many sections (**Fig. 3A**).

We first evaluated the quality of MERFISH Cellbound 1/2/3 staining by comparing to a reference staining obtained after injection of the cell membrane reporter CAAX-mCherry mRNA. CAAX-mCherry expression resulted in characteristic patterns of cell size variations with endoderm cells larger than ectoderm cells in the tailbud embryo (**Fig. 3A**). Cellbound3 staining was similar to CAAX-mCherry showing segmentation in both ectoderm and endoderm cells with appropriate shape and size (**Fig. 3A**). We excluded Cellbound2, which segmented the small yolk droplets and nuclei, leading to abnormal cell segmentation in the endoderm. PolyT co-stained with DAPI and labelled an expanded region around the nucleus in ectoderm cells but was too weak in endoderm to be applied in our system. Use of Cellbound1 resulted in a segmentation similar to Cellbound3 in ectoderm (**Fig. S2**), however, its weak signal in endoderm cells limited its usage.

We then applied the machine learning-based tool Cellpose2 (Pachitariu and Stringer, 2022; Stringer et al., 2021) to train a customized Model 1 on Cellbound3 + DAPI staining, and re-segment the cells. With this tool we were able to segment both ectoderm and endoderm cells regardless of their heterogeneous sizes in *X. laevis* embryos at different stages (**Figs 3B; S3**) and observed fewer gaps between cells, which allowed to assign 10% more transcripts (**Fig. 3C**).

While a robust Cellbound3 staining allowed good cell segmentation, it was not always of high quality in different experiments. Aiming to find the alternative membrane staining in compensation of Cellbound3, we thus compared cell segmentation by Cellbound1/2/PolyT (**Fig. S2A**) and quantified their area intersection over union (IoU) to Cellbound3 (**Fig. S2B, C**). In the ectoderm, Cellbound1-based segmentation recognized 41.2% of cells similar to Cellbound3-based segmentation, which was higher than Cellbound2 (21.7%) and polyT (19.6%) performance. However, the endoderm cells could only be segmented by Cellbound3. Indeed, we observed that the Cellbound3 signal was higher in endoderm compared to ectoderm, whereas Cellbound1 showed the opposite staining, compensating for poor Cellbound3 in ectoderm cells. Thus, to palliate Cellbound3 staining variations observed across replicate experiments, we devised an alternative cell segmentation method integrating Cellbound3 together with Cellbound1. Because Cellpose2 accepts two channels (usually the nuclear staining and one cell membrane channel), we merged the Cellbound1 and Cellbound3 channels by summing the images prior to segmentation. In an experiment with low membrane staining quality, the addition of Cellbound3 and 1 signal improved the segmentation performance compared to single channels (**Fig. S2D**). The inclusion of Cellbound1 staining signal improved the segmentation of ectoderm cells while it preserved the superior performance of Cellbound3 in endodermal regions.

In summary, we provide here a robust custom-trained Cellpose2 Model 1 for embryonic cell segmentation, yielding a proper identification of the cells, which highly improved the levels of transcripts assignment and downstream cell type analyses.

### Early-stage embryos require further optimization to identify specific cell types

Early-stage embryos such as blastulas and gastrulas represented an additional challenge for cell segmentation: our Xenopus-trained Model 1 exhibited reduced accuracy notably due to larger endoderm cells and their relatively lower staining intensity (**Fig. 4A, B**). To capture cell morphology and histology of early embryos, we custom-trained Cellpose2 Model 2 by using embryo images from earlier developmental stages, including both ectoderm and endoderm tissues. The application of the Model 2 to the MERFISH dataset from early embryos (blastula and gastrula stages) substantially improved transcript assignment efficiency to segment individual cells, reaching from 66.02% of assigned transcripts with Model 1 to 90.87% transcripts with Model 2 (**Fig. 4B, S4A**). We then compared the number of cells detected by the two models in a cell type-specific manner, using *sox17* transcripts expression as a marker for endodermal cells and *sox3* expression for ectodermal cells (**Fig. 4C**). Model 2 detected approximately three times more *sox17*-positive (endodermal) cells per embryo than Model 1, whereas both models detected a similar number of *sox3*-positive (ectodermal) cells (**Figs 4D, S4B, Table S1)**. When comparing cell size distributions, Models 1 and 2 performed similarly for small to medium-sized cells, but Model 2 showed more accuracy in detecting larger endoderm cells (**Figs 4E, S4C)**. Overlay of Model 1 and Model 2 segmentation at a later stage (tailbud St 22), with a good Cellbound staining indicated similar performances (**Fig. 4F**). These results highlight that segmentation performance in early embryos is strongly cell type- and stage-dependent. Targeted training of a cell segmentation model on challenging histology, tissue integrity and cell-type heterogeneity proved critical for achieving accurate cell boundary delineation. This approach further allowed us to maximize transcript recovery in ST analyses of a broader range of histology.

### Cell identity clustering with sub-tissue transcripts resolution at gastrula and tailbud stages

Our custom MERFISH *xenopus* gene panel contained probes for 157 genes, along with 23 non-specific (blank) probes. In total, we detected 74,178,869 transcripts across 141,059 cells from 130 sections, derived from 57 embryos and spanning three and a half MERSCOPE v1 slides.

To generate a spatially resolved reference atlas of gene expression during embryogenesis in *Xenopus laevis*, we analyzed 54 key developmental markers at gastrula and tailbud stages (st12 and 22 respectively) (Table S2, **Fig. S5A, B** raw transcripts distribution, **Fig. S6A, B** normalized gene expression per cell). As an example, detection of *myod* (paraxial mesoderm, Tajbakhsh et al., 1997) and *krt12.4* (non-neural ectoderm, Plouhinec et al., 2017) exhibited characteristic spatial distributions and expression changes with developmental stage, consistent with previous knowledge (**Fig. 5A, B).** As another example, we detected the BMP target and pluripotency factor *ventx2.1* (Bier and De Robertis, 2015; Scerbo and Monsoro-Burq, 2020) at gastrula stages, the corresponding signal vanished by late tailbud when dorsal-ventral patterning is already established. In contrast, the anteroposterior patterning ectodermal factor *pitx1,* expressed in the cement gland tissue at the anterior of the embryo (Chang et al., 2001), was only detected at the tailbud stage (**Figs S5A, B; S6A, B).** In the latter stage, we also captured increased expression of transcripts characteristic of dorsal ectoderm patterning, such as *sox8/9/10* for neural crest and *eya2/six1* for placodes. Neural ectoderm *sox2+/3+* and endoderm *sox17a+,* analyzed in Fig. 4 at earlier stages, are also shown (**Fig. S5A, B; S6A, B;** Table S2).

We noted that as for any *in situ* hybridization technique (Rademacher et al., 2025), variations in the hybridization quality for gene-specific probe sets can affect detection levels and subsequently introduce a bias in quantitative analysis. To evaluate probe efficiency, we used background signal and specificity of detection. We estimated background signal from single-transcript events (i.e., when only one gene transcript is detected in a cell, Table S2). To assess probe specificity, we calculated Moran’s I for each probe in sections from 5 embryos at gastrula stage 12 and 4 embryos from tailbud stage 22. Moran’s I ranges from −1 to 1 and reflects spatial autocorrelation, i.e. the degree of spatial clustering versus dispersion. Values close to 1 indicate spatial clustering, values close to 0 indicate spatially random expression (i.e. no spatial structure), and negative values indicate spatial anti-correlation (i.e. neighboring cells tend to have dissimilar expression levels). Most probes showed strong spatial clustering, with Moran’s I values above 0.25 as seen for *krt12.4* and *myod*. Importantly, we detected factors with tissue-restricted expression pattern such as *egr2*, *mafb* or *rax* in some embryo sections only, further highlighting the resolution and probe specificity in our datasets. Consequently, those 3 probes resulted in an overall low Moran’s I score when several sections were grouped for analysis. Last, the *tbxt* (early mesoderm marker) probe showed low detection efficiency, despite accurate *tbxt* transcripts detection by FISH in our samples (**Fig. 2A**) and was thus omitted from subsequent analysis (**Table S2 C**).

To gain insight into transcripts co-occurrence in cell types and define cell identity clusters across developmental progression, we applied Leiden clustering to data from gastrula and tailbud stages (**Figs 5C, S7**). Spatial tissue-specific organization of key marker genes expression was consistent with known histology references (**Figs 5A, B, S5, S6**). Indeed, based on detection of characteristic markers, we could resolve germ layers at gastrulation stage, i.e. *sox17a+* endoderm (clusters (c) c0, c13, c15), ectoderm into *sox2+, sox3+* neural ectoderm (c2), and *gata2+, ktr12.4+* non-neural ectoderm (c1). Neural border progenitors formed a separate *pax3+, zic1+* cluster (c8). For mesoderm, we distinguished between *chrd+, nog+* axial mesoderm (c7), *myod*+ dorsal paraxial mesoderm (c3, c4, c5, c11), *pax3+, snai1+* intermediate mesoderm (c9) and *gata2+, msx1+* ventral mesoderm (c6) (**Figs 5C, S7B**). We further identified several finer sub-tissue clusters with clear molecular signatures and co-expression of combinations of markers within the individual cells. For example, at the gastrula stage we could segregate mesoderm sub-clusters (c3, 4, 5, 6, 7, 11), based on *myod* co-expression (Chanoine and Hardy, 2003; Hopwood et al., 1992) with different markers. Within *myod*-positive mesoderm clusters, we segregated *myod+*/*epha4+/foxc2+* paraxial mesoderm I (PMI, c5, c11) and *myod+*/*epha4+/foxc2-* paraxial mesoderm II (PMII, c3, c4) based on their *foxc2* expression (Wilm et al., 2004). This is consistent with *epha4* and *foxc2* functions in fish (Durbin et al., 1998) and mice (Kume et al., 2001) and with the described genetic links (Omoteyama et al., 2007). Paraxial mesoderm II (c3, c4; *epha4+/epha2+/myod+*) shared common Ephrin signaling with both neuroectoderm (c1; *epha+*) and paraxial mesoderm I (c5, c11; *epha4+/myod+*) which is particularly interesting, given its spatial location in between these two cell layers (**Fig. 5C** top right panel; **S7B**). Finally, *myod*-negative mesoderm, including *myod-/chrd+*/*nog+* axial mesoderm (c7) and *myod-/gata2+/msx1+* ventral mesoderm (c6) were annotated based on co-expression of these other characteristic markers (Jones et al., 1995; Kelley et al., 1994). Considering Pax genes at this stage, only *pax3* was expressed, indicating the future ectoderm-derived placode and neural crest cells, and preceding *pax7* expression as previously described (Maczkowiak et al., 2010). In summary, at gastrula stage, we could assign a cell identity to 10 out of the 16 cell clusters identified with the customized Cellpose2 segmentation. This covered the diversity of tissue layers commonly identified at this stage of development. In addition, the ST clustering obtained with our 54 gene list identifies several cell subtypes yet to be fully characterized functionally (e.g. mesoderm subclusters).

At the tailbud stage (stage 22) we could assign cell identity to 24 out of 25 cell clusters found with the customized Cellpose2 segmentation, documenting the increased complexity of cell types at this early organogenesis stage compared to the gastrula stage (**Figs 5C**, middle bottom panel**; S7B**). One small cluster contained low quality cells that were not annotated (c24). Not all cell types were located on each section (e.g. eye or cement gland tissues were found on different sections) and the section shown in **Fig. 5C** exemplifies 22 clusters. In the ectoderm, *krt12.4+/cdh1+/tp63-* outer non-neural ectoderm (NNE, c0) and *krt12.4+/cdh1+/tp63+* deep inner NNE (c7) were distinct (Plouhinec et al., 2017). Three cement gland primordium subpopulations were distinguished, including a *pitx1+/otx2+/six1+* inner layer (c10), a *pitx1+/cdh1+/tp63+* intermediate layer (c19), and a *pitx1+/dlx2+/dlx3+* outer epithelium (c20) (Chang et al., 2001; Schlosser and Ahrens, 2004). Additionally, several preplacodal and placodal clusters were found (c3, c4, c5, c6, c16, c17, c23, c25) and identified by expression of *six1* or *eya1* (c4, c5, c17), or of *sox3 (*c3, *sox3+/ephA4+/gbx2+* anteroventral lateral line placode), *otx2+/pax6+* (c6, c25, optic vesicle and lens placode)*, sox9+/sox10+* (c23, otic placode) or *ephA2+/snai1+/foxc1+* early placode ectoderm tissue (c16), combined with expression of other known regional markers and their anatomic position (see gene expression details in **Fig. S7B**, Schlosser and Ahrens, 2004). Neural crest clusters comprised *snai2+/sox8+/sox10+* cells exiting the neural tube (c21), *snai1+/snai2+* early migratory neural crest cells (c12) at various anterior-posterior levels according to the section (indicated by their *hox* code), and *dlx2+/tfap2a+/sox8+* late migratory cells in the branchial arches (Kotov et al., 2024). The neural ectoderm cells were found distributed in *sox2+/sox3+/elaV+* neural tube (c9, at various anterior-posterior levels) and *chd1+/sox2+/sox3+* ventral neural tube/brain (c14). Interestingly in the mesoderm, we identified the *chd1+* notochord (c22) and three subpopulations among *myod+* myotome cells: *myod+/actc1+/zic1-* ventral somite (c2) and *myod+/actc1+/zic1+* dorsal somite (c8) and a few *actc1*+/*pax3*+/*foxc2*+ dermomyotome cells (c11) (Gaspera et al., 2012). In addition, an *hnf1b+/ephA4+* intermediate mesoderm (pronephros) population was found on some sections (c13) (Buisson et al., 2015). Last, in the endoderm, we distinguished the *pitx1+/hnfb1+/sox2+* gastrocoel roof (c15) and the *sox17+* other endoderm cells (c1). In contrast, clustering based on the default segmentation only coarsely separated mesoderm and ectoderm without resolving subclusters and failed to recover endodermal cells (**Fig. 5C**, left panel). Therefore, cell segmentation with our custom-trained models proved essential for achieving reliable cell type annotation and to assign transcripts co-expression.

In sum, these results not only provide a protocol for ST in *Xenopus laevis* embryos but also provide a ST reference gene atlas for *Xenopus*, with a combination of probes allowing to spatially resolve cell identities, and ready for further co-expression analysis of other developmental markers or genes of interest from gastrula to tailbud stages. This spatial molecular map is effective to evaluate the relative variation in gene expression levels during development, as well as to detect subtle gene co-expressions within tissues, with a single-cell resolution relying upon an accurate cell segmentation.

## Discussion

In this work, we have built a spatial map at a cellular resolution covering the dynamics of expression corresponding to key developmental markers across embryonic stages in the African clawed frog *Xenopus laevis*. We have used a MERFISH imaging-based spatial transcriptomics approach applied to Xenopus embryos. We provide a complete workflow including tissue preparation, a custom gene panel and post-imaging analysis, from cell segmentation to transcript assignment and cell identity clustering. Our method overcomes major difficulties that were constraining the application of spatial transcriptomics in the *Xenopus* community, including preparation of thin FFPE sections with good quality histology, multiplexed embedding and mounting of small embryos, and an accurate segmentation of cells. A step-by-step protocol for sample preparation and data analysis is provided as a supplementary material. Cell segmentation is key for a proper assignment of transcripts to individual cells, but often challenging in spatial transcriptomics technology, especially when the staining quality of cell membranes is poor (Petukhov et al., 2022; Yue et al., 2023). For image-based spatial transcriptomics, the current cell segmentation methods are staining-based, transcript-based, and methods combining both (Defard et al., 2026). Transcript-based methods, such as Baysor (Petukhov et al., 2022), Proseg (Jones et al., 2025), SCS (Chen et al., 2023), Comseg (Defard et al., 2024), use the spatial distribution of transcripts, optionally combined with information from nuclei or membrane markers, to assign individual RNA molecules to cells. These methods have limitations in *Xenopus*, since tools such as Comseg and SCS require nuclear staining and high transcripts density. Indeed, in embryonic cells with a size exceeding the thickness of the tissue section, nuclei are not always present. This applies to some mesoderm cells and to most endoderm cells. Thus, transcripts in cell sections without nucleus would be lost. Furthermore, lower transcript density in those germ layers, due to fewer genes included in this gene list, was not compatible with transcript-based segmentation of cells. We thus focused on staining-based segmentation and trained two Cellpose2 models on *X. laevis* embryo sections from several developmental stages, exhibiting distinct and specific histological characteristics. In this way, we could segment embryos from blastula to tailbud (stage 9 to stage 22). Both segmentation models are compatible with Vizgen postprocessing tool and other tools for spatial transcriptomics analysis workflow, such as SOPA (Blampey et al., 2024) and SpatialData (Marconato et al., 2024). Batch-effects between experiments were corrected by harmonypy (Korsunsky et al., 2019).

Improved cell segmentation allowed us to map genes co-expressed in individual cells for a refined analysis in developing tissues over time. We identified 16 gene clusters in 8 main cell types at gastrulation and 24 gene clusters subdividing 10 main tissue types at tailbud stage. Among those, we finely mapped distinct mesoderm regions in gastrula and multiple groups of ectoderm, placode, neural crest, and mesoderm cells in tailbud, based on co-expression of key factors. The provided atlas of developmental gene expression patterns over time should stimulate discovery rate in co-expression and cell identity clustering analysis. Our work demonstrates the high resolution of a semi-quantitative method for gene expression with spatial information at a single cell resolution. In the future, our gene list would benefit to extend its scope to housekeeping or cell cycle-related genes to serve as a reference for quality control and batch effect correction, and/or to assess proliferation and differentiation rates.

We believe that our workflow not only applies to *Xenopus* species but also represents an advantageous strategy for sample preparation and cell segmentation for other spatial sequencing technologies and other model organisms for developmental biology studies and pathologies where similar challenges are encountered due to complex or heterogenous nature of tissues and nuclei properties. Thus, we provide not only a developmental gene expression maps across embryogenesis in Xenopus, as a valuable resource for the community but we also pave the way for others to develop similar maps in a broad variety of models.

## Materials and Methods

### Custom gene panel design

For this atlas, 54 genes were selected based on literature knowledge of their expression and function in patterning and signaling (**Table S2**). For each gene, choice involved its expression pattern in developing cell populations, its expression levels and mRNA length, with a focus on dorsal mesoderm and ectoderm dorsal development. As *X. laevis* is pseudo-tetraploid, L or s homeolog was chosen based on the highest expression throughout stage 9 to stage 22 (Session et al., 2016). Primer sets were designed based on those informations by Vizgen and 23 non-specific primer sets (“blanks”) were added as negative controls.

### 3D mold design and printing

The embryo holder mold was designed using CAD freeware (http://www.tinkercad.com), with large cavities to optimize paraffin flow during embedding and minimize bubble formation. The design was saved in “stl” file format for subsequent 3D printing (Figs 2A, S10; Supplemental Material). Dental LT clear resin V2 (Formlabs 358-9816) was used on a Form2 3D printer (FormLabs) with a theoretical accuracy of 0.05–0.15 mm.

### Embryo injection, collection and cell membrane imaging

Animal use followed recommendations of the European Community (2010/63/UE), international guidelines and institutional Animal Care and Use Committee of Institut Curie (Authorization APAFiS#36928-2022042212033387-v1, APAFiS#964-2023092615315970 v1). For spatial transcriptomics, wild-type Xenopus *laevis* and Xenopus *tropicalis* embryos were fixed at the desired developmental stages (Faber and Nieuwkoop, 1994). From neurula stage 14 onwards, the vitelline membrane was removed manually using fine forceps in 1/3 MMR prior to fixation. For cell membrane labelling, X. *laevis* embryos were injected at 2-cell stage with CAAX-mCherry mRNA (kind gift from Dr. Lance Davidson). At the desired stage, they were fixed in formaldehyde 3.7% in PBS 1X overnight at 4°C, mounted in low-melting agarose (Sigma A9414), and either sliced to 100μm-thick sections by vibratome sectioning or simply manually bisected. After DAPI staining, they were mounted in PBS under coverslips. Images were taken with either Leica SP8 LIGHTNING Confocal Microscope or Nikon AX-R Confocal Microscope at 20x and analyzed by FIJI.

### Embedding, sectioning and mounting in paraffin

The embryos were fixed in 3.7% formaldehyde at 4 ℃ for at least overnight and up to 2 days. To maintain osmotic pressure and preserve the shape of blastocoele or archenteron, the fixative was diluted either in 1/3 MMR for X*. laevis* or in 1/9 MMR for X. *tropicalis*. After fixation, the embryos were washed in PBS, progressively dehydrated in methanol/PBS (70% methanol/PBS overnight, and 100% methanol for 3 hours) then cleared in 100% isopropanol and embedded into paraffin at 65 ℃ for storage (see details in Supplementary protocol). Before sectioning, paraffin blocks were melted and embryos were carefully oriented in fresh paraffin maintained at 70 ℃, either on top of the 3D-printed mold or at the bottom of a silicon mold, using RNAse-free dissection tools (Fig. S8). The embryos in paraffin blocks were then cut into 7μm-thick sections in the desired orientation. Sections were closely trimmed and mounted onto the MERSCOPE round coverslip, which fully dried and stored wrapped at −20 ℃ before use (Supplementary protocol).

### MERSCOPE imaging

Standard sample prep for FFPE tissues was performed following the Vizgen instruction manual, with small adaptations. In short, a cell boundary staining was applied, followed by hybridization with the custom gene panel. Then the section was embedded and cleared. For extended clearing of the highly autofluorescent frog tissue (resistant FFPE sample), digestion was prolonged to 4h at 37 °C, followed by clearing overnight at 47°C. Right before imaging, the section was embedded again, placed under a weight (e.g. the lid of 50 ml Falcon tube filled with 2 ml RNAase-free water) to avoid tissue lifting during acquisition. Slides were imaged with the Vizgen MERSCOPE microscope and processed with the software version 233.

### Customized Cellpose2 models training and cell re-segmentation

For training the custom Cellpose2 Models 1 and 2, images were cropped from section stainings using the Cellbound3 and DAPI channels. To train Model 1, a gastrula-stage 12.5 section and a tailbud-stage 22 section were used. For Model 2, we used two sections per stage from blastula-stage 9 and gastrula-stages 10.5, 11 and 12 embryos. These sections were cropped from the MERSCOPE Raw Cellbound images into smaller tiles. Cells in each tile were manually annotated and then used to train Cellpose2. Cell re-segmentation and generation of a new transcripts matrix and cell metadata were done in the Vizgen post-processing tool (VPT) version 1.3 with the Cellpose option using our customized Cellpose2 models – Model 1 and Model 2. While applying Model 1, we used cellprob_threshold = 0 and flow_threshold = 0.4 as parameters in the VPT. Similarly for Model 2, we used cellprob_threshold = 0 and flow_threshold = 0.95. Furthermore, for Model2, cells with a total transcripts number inferior to 10 were excluded to remove false positive cells.

### Imaging pre-processing

To merge Cellbound1 and Cellbound3 stainings, we first normalized their intensity to an equivalent level. The mean intensity of background or signals from the 5 representative selections was measured, and the ratio was calculated as the mean intensity of Cellbound1 divided by that of Cellbound3. For signal regions, the Cellbound1-to-Cellbound3 ratio was 1.56. For background regions, the Cellbound1-to-Cellbound3 ratio was 1.69. Thus, we normalized the Cellbound3 by multiplying the image by 1.69. Then the Cellbound1 and normalized Cellbound3 images were merged into one image by addition or maximum. The intersection over union (IoU) of the segmented mask was calculated using the FUJI plugin MiC (https://github.com/MultimodalImagingCenter/MiC).

### Data processing, Statistics and visualization

The statistics of the probes were calculated by scanpy.pp.calculate_qc_metrics or numpy. Moran’s I was calculated by Squidpy (Palla et al., 2022). The clustering was calculated by Scanpy: we calculated the PCA and computed a neighborhood graph and UMAP, then clustered using the Leiden algorithm with resolution 1.0 (McInnes et al., 2020; Traag et al., 2019; Wolf et al., 2018).

### *In situ* hybridization

*In situ* hybridization was done as in Monsoro-Burq, (2007) with adaption for FFPE sections: the FFPE sections were deparaffinized for 20min by X-free Solvant xylene substitute (Bio-optica #06-1305F) and rehydrated through 100%, 95%, 85%, 70% ethanol washes and rinse in PBS with 0.1% Tween 20. Then the sections were rinsed with 2xSSC (Euromedex #EU0300-C) and hybridized with DIG-labeled antisense probes in 65 ℃ overnight. RNAase treatment was skipped. After colorimetric staining, the sections were stained by 0.125% of eosin (Bio-optica #05-10007/L) for 1min, then mounted in entellan (Sigma #1.07961.0500) and imaged using ZEISS Lumar equipped with an ICC color camera or using ZEISS Axio Imager 2.

## Supporting information

Combined Supplemental Materials

## DATA AVAILABILITY

Materials and codes are available upon request to the corresponding author and on Monsoro-Burq Lab github page (https://github.com/monsoro, forked from https://github.com/chenxiZ7/MERFISH_pipeline).

## ACKNOWLEDGEMENTS

We thank Dr Lance Davidson for kind gift of the CAAX-mCherry plasmid. We thank all members of the Monsoro-Burq and Almouzni teams for their support. We thank the following members of the Institut Curie core facilities: T. Herpin, J. Roustel and H. Gautier from the In vivo experimentation platform for animal care; S. Leboucher, R Leclere and A. Nicolas from the Histopathology platform for technical help for preparation of samples, C. Messaoudi, L. Besse and M.N. Soler from the Multimodal Imaging Center for technical help for imaging processing and quantification, and the Institut Curie Single Cell initiative. Funding to A.H.M.B. includes the Agence Nationale pour la Recherche (ANR-15-CE13-0012-01; ANR-21-CE13-0028 including support for CZ), Institut Universitaire de France and Fondation pour la Recherche Médicale (DEQ20150331733 and FDT202404018665 including support for CZ). Funding to G.A. laboratory includes the European Research Council (ERC-2015-ADG-694694 ‘ChromADICT’), the Ligue Nationale contre le Cancer (Equipe labellisée Ligue), France and Agence Nationale de la Recherche, France (ANR-11-LABX-0044_DEEP, ANR-10-IDEX-0001-02 PSL) and ‘Cellular identities and destinies’ exploratory research program (PEPR Cell-ID), managed by the Agence Nationale de la Recherche France 2030 program reference number ANR-24-EXCI-0001, ANR-24-EXCI-0002, ANR-24-EXCI-0003, ANR-24-EXCI-0004, ANR-24-EXCI-0005’. SD was supported by the European Union’s Horizon 2020 research and innovation program under the Marie Skłodowska-Curie grant agreement EuReCa No 847718, attributed to IS and GA.

## AUTHOR CONTRIBUTIONS

A.M.-B conceptualized the initial study; GA and IS extended the scope to the early development. C.Z. and A.M.-B. developed methodology. <u>Investigation</u>: C.Z., S.S and A. M.-B designed the gene panel; C.Z., S.D., V.K., I.S. and A.M.-B. prepared the samples; K.E.B processed slides through Vizgen apparatus. <u>Software, data curation, formal analysis</u>: T.D developed the code segmentation feature, wrote tutorials and trained C.Z.; C.Z., T.D. and S.D. processed the raw data and optimized cell segmentation. C.Z., S.D., I.S. and A.M.-B. analyzed data. <u>Visualization</u>: C.Z. and S.D. generated the figures and tables. <u>Writing</u>: C.Z., I.S, S.D., G.A. and A.M.-B. wrote and reviewed the manuscript. Critical reading and discussion of data involved all authors. <u>Supervision</u>: T.W. supervised T.D. work; A.M.-B. supervised C.Z., V.K., S.S. work. G.A. and I.S. supervised S.D. work. <u>Resources, project administration, funding</u> <u>acquisition</u>: I.S, G.A. and A.M.-B.

