## Supplementary material for "An optimized workflow for spatial transcriptomics across early development in Xenopus": Combined Supplemental Materials

#### SUPPLEMENTARY TEXT

#### 1. SUPPLEMENTARY TABLE LEGENDS

**Table S1. Summary table of cell type distribution by Model 1 and Model 2**

**Table S2. Selected genes in the reference atlas and their expression across developmental stages**

**Table S3. Evaluation of the gene panel efficiency.** We evaluated the statistics of each gene expression and assessed the probe efficiency. High-efficiency probes were defined as those with Moran's I >= 0.25 in either gastrula or tailbud stages. *hoxc10.L*, *hoxc9.s*, *hoxd11.L* and *olig4.L* exhibited medium efficiency with weak spatial specific pattern. Only *tbxt.s* showed low efficiency: because of its low detection in MERFISH but high detection by classical in situ hybridization, this probe should be discarded or re-designed. Probes for *egr2.L* and *rax.L* displayed high efficiency, as they were detected exclusively in tissue sections containing their corresponding expression tissues, and showed minimal expression in the sections in Fig. S5 and Fig. S6 which lacked those target tissues. Mean count refers to the average expression level of genes across all cells. A single-transcript event refers to cases where only one transcript of a gene is detected in a cell. Potential background was estimated by percentage of single transcript event (number of single transcript event \ total number of cell).

#### 2. SUPPLEMENTARY FIGURE LEGENDS

**Figure S1 (related to Fig.2). Hematoxylin & Eosin staining of Xenopus embryos at different developmental stages to assess the quality of histology.** (A) H&E staining of the sections used for MERFISH experiment in Fig.2 (B) from gastrula stage 9-12 Xenopus laevis embryos. Scale bar = 500µm. (B) H&E staining of the sections from blastula (St 7), gastrula (St 11) and neurula (St 18) Xenopus tropicalis embryos. Scale bar = 500 µm.

**Figure S2 (related to Fig.3). An alternative segmentation method by integrating Cellbound1 and Cellbound3, when Cellbound3 staining is weak.** (A) Using a custom Cellpose2 Model 1, cell segmentation obtained by combining Cellbound1, Cellbound2, PolyT, Addition Cellbound1+3, or Max Cellbound1/3 with DAPI (red) is overlaid on cell segmentation obtained with Cellbound3+Dapi (gray). The figure shows representative regions of the ectoderm and endoderm. (B)-(C) Intersection over Union (IoU) between the cell segmentation (red in A) with Cellbound 3 + Dapi (gray in A) indicates the quantification of segmentation overlay. (B) In ectoderm region, Cellbound1+3 addition showed 83.7% overlap compared to Cellbound3 alone; While in endoderm region, Cellbound 1/3 max showed 54.7% overlap compared to Cellbound3. (C) In whole section, Cellbound 1+3 addition showed 61.7% overlap compared to Cellbound3, performed better than Cellbound1/3 max. IoU was defined with threshold as 80%. (D) Comparison of cell segmentation based on Cellbound1, Cellbound3, PolyT, or Addition Cellbound1+3 with DAPI for Cellbound3 suboptimal section. With the compensation of two staining, the addition Cellbound1+3 with DAPI are able to segment adequately both the endoderm and ectoderm cells.

**Figure S3 (relates to Fig. 3). Model 1 provides accurate cell segmentation from stage 12.5 to stage 22 in *Xenopus laevis* embryos.**

**Figure S4 (relates to Fig.4). Comparison of Model 1 and Model 2.** (A) Percent of transcripts captured inside cell boundaries by Model 1 and Model 2. (B) Violin plot of cell volume distribution of different cell categories between Model 1 and Model 2 (arbitrary units, a.u.). Cells with *sox3* transcripts ≥ 9 were classified as ectoderm, cells with *sox17a* transcripts ≥ 5 as endoderm, cells meeting both criteria as double positive, and cells meeting neither criterion as double negative. (C) Scatter plot of endoderm cells (*sox17a* positive) as detected by Model 1 (blue) and Model 2 (yellow). Y axis represents the transcript counts of *sox17a* and X axis their corresponding cell volume (a.u.).

**Figure S5 (relates to Fig.5). Spatial distribution of the 54 probes on a transverse section of a *X. laevis* gastrula at stage 12 (A) and tailbud at stage 22 (B).** Each red dot represents a signal from a single mRNA molecule in principle. Embryo section histology is shown as cell masks. D: Dorsal, V: Ventral, scale bar = 500µm.

**Figure S6. (relates to Fig. 5). Spatial gene expression pattern in a transverse section of a *X. laevis* gastrula at stage 12 (A) and tailbud at stage 22 (B).** Expression levels for the genes were calculated by volume-normalization and log-transforming the individual transcript counts per cell, then represented by greyscale values within a cell. Embryo section morphology represented by cell masks. D: Dorsal, V: Ventral, scale bar = 500 µm.

**Figure S7. (relates to Fig. 5). Analysis of gene-based cell clusters in *X. laevis* gastrula stage 12 and tailbud stage 22.** (A) UMAP plots of gene-based clusters. Leiden clustering with resolution = 1 on cells from five embryo sections at gastrula stage 12, and four embryo sections at tailbud stage 22. (B) Dot plot representing mean expression of top 3 genes per cluster (red gradient), and percentage of cells containing these genes per cluster (circle size). Cell types were assigned based on gene expression and spatial location. At gastrula stage 12, we identified 10 main cell types for 12 gene clusters and four unidenfitied ones – Endoderm (Endo, c0, c13, c15), Neural ectoderm (NE, c1), Neural border (NB, c8), Non-neural ectoderm (NNE, c2), Axial mesoderm (AM, c7), Intermediate mesoderm (IM, c9), Paraxial mesoderm (PM Ⅰ and PM Ⅱ, c3, c4, c5, c11), and Ventral mesoderm (VM, c6). At tailbud stage 22, we identified 12 main cell types subdivided into 24 gene clusters and one unidentified one: Endoderm (Endo, c1, 15), Brain (c14), Neural tube (NT, c9), Neural crest (NC, c12, 18, 21), Placode (c3, 4, 5,16, 17, 23, 25), Non-neuroectoderm (NNE, c0, 7), Optic vesicle (OV, c6), Notochord (Noto, c22), Myotome (c2, 8), Dermomyotome (DM, c11), Intermediate mesoderm (IM, c13), and Cement gland (CG, c10, 19, 20). Clusters with low cell numbers and without a specific spatial pattern are labelled as NA.

**Figure S8.** Equipment: (A) 3D-printed platform located in the embedding mold; (B) dissection tools used to manipulate the embryos; (C) The dry heating block used to warm paraffin during the embedding. (D) Microtome.

**Figure S9.** Embedding setting: (A) the embedding silicon mold is warmed to 70°C; (B) if needed, stabilize heater; (C) the binocular stereoscope should have a long working distance; (D) RNAse-free warm dissection tools are used to orientate the embryos; (E) Bensen burner to maintain the dissection tools warm during embedding.

**Figure S10.** (A) 3D-printed platform, with hole diameters of 1.0 mm, which is adapted to embryos from stage 9 to stage 22 has been optimized for liquid paraffin circulation under the embryos. Other sizes of the 3D-printed platform may be needed for other stages. (B) The embryos are oriented manually on the 3D-printed platform under the stereoscope.

**3. STEP BY STEP SAMPLE PREPARATION PROTOCOL**
**(using the MERFISH technology, MERSCOPE, Vizgen)**

#### A. Reagents, Materials and Equipment

**1. Reagents:**

- Formaldehyde 37%, Sigma 252549
- [R] Fixative
- Isopropanol, ultrapure, Honeywell 33539
- Methanol, ultrapure, VWR, 20821.310
- [R] MMR 10× (Marc's Modified Ringer's solution 10×)
- [R] Phosphate-buffered saline (PBS 10×)
- Paraffin, melting temperature at 54 – 56 °C, VWR 2338900
- RNaseZAP, Sigma R2020
- RNase Inhibitor, Murine (NEB M0314L)
- Proteinase K, Molecular Biology Grade (NEB P8107S)
- MERSCOPE 140 Gene Panel V 2.0, 10 samples
- MERSCOPE FFPE sample prep kit 10400114 containing:

Deparaffinization buffer (PN 20300112)
Decrosslinking Buffer (PN 20300115)
Pre-Anchoring activator (PN 20300113)
Formamide wash buffer (PN 20300002)
Anchoring buffer PN 20300117)
Gel coverslip (PN 30200004)
Gel embedding Premix (PN 20300118)
Digestion Premix (PN 20300005)
Clearing Premix (PN 20300114)

- MERSCOPE Cell Boundary Stain Kit, Vizgen 10400118 containing:
- Cell Boundary Blocking Buffer premix (PN20300012)
- Cell Boundary Primary Stain mix (PN20300010)
- Cell Boundary Secondary Stain mix (PN20300011)
- MERCOPE FFPE standard slides box, with 1.25 $cm^2$ imaging area, Vizgen 10500110
- MERSCOPE large FFPE slides box, with 3 $cm^2$ imaging area, Vizgen 10500121
- MERSCOPE imaging cartridge 300 genes, custom, Vizgen 10400005

**2. Materials and Equipment (Figures S8, S9, S10):**

- 3ml plastic pipette with 2mm opening, Thermo Scientific™ 225NL-1S
- 3D-printed platform with holes to hold the embryos, custom made
- 6-well cell culture plate (any brand)
- Aluminum foil
- Bunsen burner
- Dissection needles (e.g.labbox.com DISS-001-005 and DISS-002-005)
- Dry block heater
- MERSCOPE Instrument (MERSCOPE Ultra™, Vizgen)
- MERSCOPE Photobleacher, Vizgen 10100003
- Microtome, LEICA RM2245
- Painting brushes to manipulate the paraffin ribbons and sections

- Parafilm
- Stereoscope with a long working distance, LEICA WILD MZ8
- Silicone rubber Embedding mold (25mm × 25mm × 25mm)
- Oven, set to 65 °C.
- Ventilated oven, 42°C

**3. Recipes**

---

**Fixative** (3.7% formaldehyde in 1/3 MMR), **100mL**

| | |
|---|---|
| 37% Formaldehyde | 10 ml |
| 1/3 MMR | 90 ml |

Make fresh each time

---

**MMR 10× (Marc's Modified Ringer's solution 10×), 1L**

58.44g NaCl
1.5g KCl
2.5g MgSO4
2.9g CaCl2
50ml 1M HEPES (pH 7.8)
2ml 0.5M EDTA
Dissolve in 900ml $H_2O$, adjust pH to 7.4. Adjust to 1L with Milli-Q water. Store at 4°C.

---

**1/3 MMR, 3L**

100 ml MMR 10×
2.9 L Milli-Q water

---

**10X PBS, 1L**

80g NaCL
2g KCl
14.4g $Na_2HPO_4$
2.4g $KH_2PO_4$
Dissolve in 900ml RNase-free $H_2O$ and adjust the pH to 7.4. Adjust to 1L with RNase-free $H_2O$.

---

**4. General notes**

1. All solutions are RNase-free and made using RNAse-free Millipore-purified water;
2. Rinses are done without an incubation time, washes are incubated as long as indicated;
3. All equipment and materials need to be RNase-free:
    i. Metal, plastic and glass tools are cleaned by a 30 min incubation in 20-30% bleach (made in tap water), and are rinsed 5 times with MilliQ water;
    ii. Silicone embedding molds, 3D-printed platform, brushes are cleaned with 1% dish washing soap for 30 min and rinsed 5 times with MilliQ water.
    iii. Cleaned items may be air dried, wrapped and stored in clean aluminum foil;
    iv. The bench and dry block heater are cleaned with detergent and covered by an aluminum foil.
    v. Microtome and stereoscope are cleaned by RNaseZAP
4. Paraffin should be melted overnight at 65°C in advance of usage.

**5. Custom gene panel design**

see main Materials and Methods

#### B. Embryo FFPE sample preparation

##### 1. Xenopus embryo/tadpole collection (2-10 days)

Embryos are collected at the desired developmental stage according to standard protocols (Faber and Nieuwkoop, 1994; Sive et al., 2007) and following national and international guidelines on animal welfare (see: https://www.xenbase.org/xenbase/anatomy/alldev.do).

##### 2. FFPE sample preparation (3 days)

The FFPE sample preparation protocol has been validated for *X. laevis and X. tropicalis* embryos.

2.1. Collect the embryos at the desired stages, open the vitelline envelope if needed:
*Opening the vitelline membrane or not does not influence the quality of the experiment but optimizes the morphology of neurula-stage and tailbud/tadpole stage embryos.*

2.2. Fixation of the embryos:

*i.* Transfer the embryos in fixative at 4°C overnight with gentle nutation *(use the lowest speed, e.g. 7-10 rpm. A maximum of 2 days in fixative at 4 ℃ has been validated.*)

ii. Wash 3 times for 10 min in 1×PBS at room temperature.

2.3. Dehydration (3 hours):

*From this step on, embryos are fragile, each medium change should be done avoiding touching them.*

i. Dehydration in 50% methanol and 50% PBS for 30 min at room temperature;

ii. Dehydration in 70% methanol and 30% PBS overnight at 4°C;

iii. Dehydration in 100% methanol with 3 washes for 20 min each at room temperature, with gentle nutation.

*iv.* Dehydration in 100% methanol for 2 hours at 4°C, with gentle nutation. *Do not leave embryos in 100% methanol overnight for paraffin embedding purposes; it makes the embryos brittle and affect the quality of histology.*

2.4. Methanol removal and replacement by isopropanol (3 hours):

i. Wash 3 times in 100% isopropanol for 20min each at room temperature with gentle nutation;

ii. Wash 2 hours in 100% isopropanol at room temperature with gentle nutation;

iii. with minimum speed on the rotator (about 10 rpm).

*Methanol is poorly miscible with paraffin, whereas isopropanol is miscible with both paraffin and methanol. Replacement of methanol by isopropanol in embryos optimizes homogeneous penetration of paraffin into tissues and cavities.*

2.5. Paraffin inclusion

i. Paraffin bath 1: replace isopropanol by liquid (65°C) paraffin and incubate for 1 hour at 65°C.

ii. Paraffin bath 2: change to fresh liquid (65°C) paraffin and incubate overnight at 65°C.

iii. (optional) Embed the embryos without orienting them in fresh paraffin and cool down at RT.

*If the embryos are not to be oriented and embedded immediately after paraffin bath 2, embryos can be stored in paraffin after 2 additional one hours of paraffin baths and stored at room temperature. Embryos can be stored in 6/24-well plates or aliquoted in small silicone embedding mold (Sigma embedding molds E-4390). Before next step, melt the embryos overnight at 65 ℃.*

##### 3. Orientation of multiple embryos in paraffin (1.5 hour)

*We provide an optimized method by introducing continuous heating in a dry heater and 3D-printed platform under stereoscope, allowing to embed up to 40-50 embryos together.*

3.1. Pre-warm all embedding tools at 65°C, 2 hours before embedding (silicon paraffin mold, 3D-printed mold, 3ml plastic pipette, and dissecting needles). Stably install equipments as shown in Fig. S9 as any vibration of the bench would affect embryo orientation.

i. Under the stereoscope, set the dry heater to 70°C to maintain paraffin liquid.

ii. Install the 3D-printed platform in the silicon mold filled with freshly melted paraffin. Remove all bubbles and transfer the silicon mold with the platform into the dry heater.

3.2. Transfer all embryos onto the side of the 3D-printed platform using a pre-warmed 3ml plastic pipette.
3.3. Orient the embryos as needed using pre-warmed or flame-heated dissecting needles (**Fig. S9**).
*3.4.* Turn off the dry bath, let the paraffin solidify. *Do not move the silicon mold until paraffin solidifies (about 1 hour at RT) to avoid altering embryo orientation. To finish solidifying the paraffin, cool the silicon mold on ice for at least 2 hours, or overnight at room temperature.*

**4. Sectioning and mounting on the MERSCOPE slide (1 day)**

4.1. Clean the microtome and all histology instruments (heating plate, painting brush, etc) with RNaseZAP. Wear a surgical mask and tied hair to avoid RNAse contamination from the experimenter.
*4.2.* Closely trim the paraffin block and cut 7µm-thick sections; collect them on a clean paper surface with the painting brushes; *(alternatively, larger sections can be cut and trimmed afterwards).*
4.3. Place the MERSCOPE slide on a heating plate kept at room temperature during mounting.
   *i.* Draw a hydrophobic circle along the marked line of imaging region using paraffin. *A "paraffin pen" can be home-made by immersing a dissection needle into melted paraffin and let it solidify.*
   ii. On the paper sheet, closely trim the extra paraffin around sections using a fresh razor blade.
   iii. With a pipetman, add small drops of room-temperature water at the desired position of the MERSCOPE slide and transfer one section on top of it with dissection needles. Add water to exactly underlay the section. Too much water would let the section drift out of place. Use a clean painting brush to gently guide the sections if needed.
   iv. To embed multiple sections, repeat until all the sections are mounted on the slide.
   v. Add water to completely fill space underneath all sections, avoiding spilling outside the marked imaging region, which would cause section drifting.
4.4. Turn on the heating plate at 42°C, wait until all sections are flat, all wrinkles expanded. Then gently absorb the water using tissue paper put at a corner without sections.
4.5. Let the slides fully dry in a ventilated oven at 42°C overnight. The Merscope slides should be stored dried at -20°C in a parafilm-sealed RNAse-free container before further processing, for a maximum of 1 month. *Do not risk breaching the edge of the slide using forceps as this would cause slide breaking during the imaging process. Use clean gloves and hand manipulation instead.*

#### C. MERSCOPE slide preparation (5 days)

The samples were prepped following the Vizgen instruction manual for FFPE tissues, with minor adaptations (marked below in *italic)*.

**5. Deparafinization and Decrosslinking**

5.1. Ensure the sample is completely dry before proceeding.
5.2. Add **500 µL** Deparaffinization Buffer (PN 20300112) onto the center of the FFPE tissue section. Ensure the Deparaffinization Buffer covers the whole tissue section and try to prevent the Deparaffinization Buffer from flowing beneath the MERSCOPE FFPE Slide. Incubate at 55°C (oven) for 5 min.
5.3. Aspirate the Deparaffinization Buffer and repeat step 2.
5.4. Aspirate the Deparaffinization Buffer and add **500 µL** Deparaffinization Buffer onto the center of the tissue section. Ensure the Deparaffinization Buffer covers the whole tissue section and try to prevent the Deparaffinization Buffer from flowing beneath the MERSCOPE FFPE Slide. Incubate at room temperature for 5 min.
5.5. Aspirate the Deparaffinization Buffer. Wash **3x** with **5 mL** 100% ethanol, incubate 2 min each wash. If Deparaffinization Buffer is underneath the MERSCOPE FFPE Slide, gently lift the slide to ensure oil droplets (resulting from deparaffination) from underneath are also removed. Transfer to a new labeled petri dish if this facilitates oil droplet removal.

5.6. **IF** oil droplets are observed after 3x washes, repeat 100% ethanol washes until no oil droplets are observed.
5.7. Wash **1x** with **5 mL** 90% ethanol, incubate 2 min.
5.8. Wash **1x** with **5 mL** 70% ethanol, incubate 2 min.
5.9. Aspirate the 70% ethanol *and air dry for 30min at RT*
5.10. Add **5 mL** Decrosslinking Buffer (PN 20300115).
5.11. Aspirate and add **5 mL** Decrosslinking Buffer. Place the petri dish onto a heating pan and incubate at 90°C (oven) for 15 min. **DO NOT** heat above 95°C.
5.12. The petri dish is hot and should be handled accordingly: Remove the petri dish from the oven and cool on the bench for 5 min.

**6. Anchoring Pretreatment**

6.1. Aspirate the Decrosslinking Buffer.
6.2. Wash 2x with 5 mL Conditioning Buffer (PN 20300116), incubate 1 min each wash.
6.3. Add 5 mL Conditioning Buffer, incubate at 37°C for 30 min in an incubator.
6.4. Prepare Pre-Anchoring Reaction Buffer:

| Pre-Anchoring Reaction Buffer | 1 sample |
|---|---|
| Conditioning Buffer (PN 20300116) | 100 µL |
| Pre-Anchoring Activator (PN 20300113) | 5 µL |
| RNase inhibitor | 5 µL |

6.5.
6.6. First aspirate the Conditioning Buffer to dry the region of MERSCOPE FFPE Slide that does not have tissue section. Then carefully aspirate around the tissue section to remove extra solution without touching the tissue section. The tissue section should not be completely dry for more than 1 min.
6.7. Add 100 µL Pre-Anchoring Reaction Buffer onto the tissue to cover the whole tissue. Use scissors to cut a piece of parafilm 2×2 cm. Use tweezers to peel off the parafilm backing and place the side previously protected by the backing onto the solution. Avoid introducing air bubbles.
6.8. Seal the petri dish with parafilm and place in a humidified 37°C cell culture incubator for 2h.

**7. Cell Boundary Staining**

7.1. Use tweezers to remove the parafilm. Add 5 mL 1X PBS, incubate 2 min.
7.2. Prepare Blocking Solution:

| Blocking Solution | 1 sample |
|---|---|
| Blocking Buffer C Premix (PN 20300100)<br>**OR**<br>Cell Boundary Blocking Buffer Premix (PN 20300012) | 100 µL |
| RNase inhibitor | 5 µL |

7.3. Aspirate the 1X PBS to dry the MERSCOPE FFPE Slide, leaving just enough liquid to cover the tissue section.
7.4. Add **100 µL** Blocking Solution onto the center of the tissue section. Use scissors to cut a piece of parafilm 2×2 cm. Use tweezers to peel off the parafilm backing and place the side previously protected by the backing onto the solution. Avoid introducing air bubbles.
7.5. Incubate at room temperature for 1 h.
7.6. Prepare Primary Staining Solution:

| Primary Staining Solution | 1 sample |
|---|---|
| Blocking Buffer C Premix (PN 20300100)<br>**OR**<br>Cell Boundary Blocking Buffer Premix (PN 20300012) | 100 µL |
| RNase inhibitor | 5 µL |
| Cell Boundary Primary Stain Mix (PN 20300010) | 1 µL |

7.7. Use tweezers to remove the parafilm.

7.8. Aspirate the solution to dry the MERSCOPE FFPE Slide, leaving just enough liquid to cover the tissue section.

7.9. Add **100 µL** Primary Staining Solution onto the center of the tissue section. Use scissors to cut a piece of parafilm 2×2 cm. Use tweezers to peel off the parafilm backing and place the side previously protected by the backing onto the solution. Avoid introducing air bubbles.

7.10. Incubate at room temperature for 1 h.

7.11. Use tweezers to remove the parafilm.

7.12. Wash **3x** with **5 mL** 1X PBS, incubate 5 min on a rocker each wash.

7.13. Prepare Secondary Staining Solution:

| Secondary Staining Solution | 1 sample |
|---|---|
| Blocking Buffer C Premix (PN 20300100)<br>**OR**<br>Cell Boundary Blocking Buffer Premix (PN 20300012) | 100 µL |
| RNase inhibitor | 5 µL |
| Cell Boundary Secondary Stain Mix (PN 20300011) | 3 µL |

7.14. Aspirate the 1X PBS to dry the MERSCOPE FFPE Slide, leaving just enough liquid to cover the tissue section.

7.15. Add **100 µL** Secondary Staining Solution onto the center of the tissue section. Use scissors to cut a piece of parafilm 2×2 cm. Use tweezers to peel off the parafilm backing and place the side previously protected by the backing onto the solution. Avoid introducing air bubbles.

7.16. Incubate at room temperature for 1 h.

7.17. Use tweezers to remove the parafilm.

7.18. Wash **3x** with **5 mL** 1X PBS, incubate 5 min on a rocker each wash.

7.19. Aspirate the 1X PBS. In a fume hood, add **5 mL** fixation buffer to fix the stained tissue section at room temperature for 15 min.

7.20. Wash **2x** with **5 mL** 1X PBS, incubate 5 min each wash.

7.21. Proceed immediately to the next step.

**8. RNA Anchoring**

8.1. Aspirate the 1X PBS (if continuing from cell boundary staining) or discard parafilm (if continuing from anchoring pretreatment) and wash 1x with 5 mL Sample Prep Wash Buffer (PN 20300001).

8.2. Add 5 mL Formamide Wash Buffer (PN 20300002), incubate at 37°C for 30 min in an incubator in a fume hood.

8.3. Aspirate the Formamide Wash Buffer to dry the region of MERSCOPE FFPE Slide that does not have the tissue section. Then carefully aspirate around the tissue section to remove extra Formamide Wash Buffer without touching the tissue section. The tissue section should not be completely dry for more than 1 min.

8.4. Add 100 µL Anchoring Buffer (PN 20300117) onto the center of the tissue section. Use scissors to cut a piece of parafilm 2×2 cm. Use tweezers to peel off the parafilm backing and place the side previously protected by the backing onto the solution. Avoid introducing air bubbles. Seal the petri dish with parafilm and place in a humidified 37°C cell culture incubator overnight (12-18 h).

8.5. Following overnight incubation, use tweezers to remove the parafilm and add **5 mL** Formamide Wash Buffer.

8.6. Incubate at 47°C for 15 min in an incubator in a fume hood.

8.7. Wash **1x** with **5 mL** Sample Prep Wash Buffer, incubate 2 min.

8.8. Proceed immediately to the next step.

**9. Gel Embedding**

9.1. Clean a Gel Coverslip (PN 30200004) by spraying with RNaseZap solution and wiping with a Kimwipe, followed by spraying 70% ethanol and wiping with a Kimwipe.

9.2. Add 100 µL Gel Slick Solution onto the Gel Coverslip. Allow the Gel Slick Solution to evaporate for 10 min at room temperature. Wipe gently with a Kimwipe to remove any remaining film, liquid, or deposition from the glass. Use immediately after preparation.

9.3. Prepare Gel Embedding Solution:

| Gel Embedding Solution | 1 sample |
|---|---|
| Gel Embedding Premix (PN 20300118) | 5 mL |
| 10% w/v ammonium persulfate solution | 25 µL |
| N,N,N′,N'-tetramethylethylenediamine | 2.5 µL |

9.4. Aspirate the Sample Prep Wash Buffer. Retain 100 µL Gel Embedding Solution in a small tube. Add the remainder of the 5 mL Gel Embedding Solution to the sample, ensure the sample is fully covered, and incubate at room temperature for 1 min.

9.5. Using a pipette, transfer the majority of the Gel Embedding Solution to a waste tube (to monitor the gel formation).

9.6. Aspirate to dry the MERSCOPE FFPE Slide, leaving just enough liquid to cover the tissue section.

9.7. Add 50 µL of the retained Gel Embedding Solution on the tissue section.

9.8. Place the tips of one pair of tweezers on an area of the MERSCOPE FFPE Slide without touching the tissue section. Use tweezers to pick up the 20-mm Gel Slick-treated Gel Coverslip. With the Gel Slick-treated side facing down toward the tissue, place the edge of the Gel Coverslip against the tweezer tips resting on the MERSCOPE FFPE Slide, creating stability, and slowly lower the Gel Coverslip onto the tissue section to spread the Gel Embedding Solution. If needed, adjust the Gel Coverslip so it is positioned in the center of the MERSCOPE FFPE Slide. Gently press the Gel Coverslip to squeeze out excess Gel Embedding Solution and remove the extra Gel Embedding Solution by aspiration.

9.9. Incubate at room temperature for 1.5 h.

9.10. Gently brace the Gel Coverslip with tweezers in one hand and lift the 20-mm Gel Slick-treated Gel Coverslip with the sharp tip of a Hobby Blade and discard the Gel Coverslip appropriately.

**10. Clearing – Resistant FFPE Tissue**

10.1. Prepare Digestion Mix:

| Digestion Mix | 1 sample |
|---|---|
| Digestion Premix (PN 20300005) | 200 µL |
| RNase inhibitor | 5 µL |

10.2. Aspirate to dry the MERSCOPE FFPE Slide without touching the gel. Add 200 µL Digestion Mix onto the gel. Use scissors to cut a piece of parafilm 2×2 cm. Use tweezers to peel off the parafilm backing and place the side previously protected by the backing onto the solution. Avoid introducing air bubbles.

10.3. Incubate at 37°C *for 4 hours.*

10.4. Warm Clearing Premix (PN 20300114) at 37°C for 30 min before use. The Clearing Premix should be a clear solution before use. If the solution is cloudy, warm until the solution becomes clear. Prepare Clearing Solution:

| Clearing Solution | 1 sample |
|---|---|
| Clearing Premix (PN 20300114) | 5 mL |
| Proteinase K | 50 µL |

10.5. Aspirate the Digestion Mix. Add **5 mL** Clearing Solution.
10.6. Place the lid on the petri dish and spray the outside with 70% ethanol to sterilize.
10.7. Seal the petri dish with parafilm and place in a humidified 47°C cell culture incubator for 24h. **DO NOT** incubate at 47°C >24 h otherwise the RNA will begin to degrade - this is important to remember if clearing over the weekend.
10.8. At this point, samples can be stored in clearing solution at 37°C up to 4 days.

**11. Autofluorescence Quenching**
11.1. Aspirate bubbles or condensation from the lid of the petri dish to minimize light scattering from above.
11.2. Place the parafilm-sealed petri dish in the MERSCOPE Photobleacher (PN 10100003). **ENSURE** there are no labels/writing/other items on the lid that may block the light. *Use only the minimum or parafilm necessary.*
11.3. Turn on the MERSCOPE Photobleacher and leave at room temperature *for 5 h.*

**12. Encoding Probe Hybridization**
12.1. Aspirate the Clearing Solution and wash 3x with 5 mL Sample Prep Wash Buffer (PN 20300001), incubate at room temperature for 5 min on a rocker each wash.
12.2. Add 5 mL Formamide Wash Buffer (PN 20300002), incubate at 37°C for 30 min in an incubator in a fume hood.
12.3. First aspirate the Formamide Wash Buffer to dry the region of MERSCOPE FFPE Slide that does not have gel. Then carefully aspirate around the gel to remove extra Formamide Wash Buffer without touching the gel. The gel should not be completely dry for more than 1 min.
12.4. Add **100 µL** MERSCOPE Gene Panel Mix onto the center of the gel. Use scissors to cut a piece of parafilm 2×2 cm. Use tweezers to peel off the parafilm backing and place the side previously protected by the backing onto the solution. Avoid introducing air bubbles.
12.5. Place the lid on the petri dish and spray the outside with 70% ethanol to sterilize.
12.6. Seal the petri dish with parafilm and place in a humidified 37°C cell culture incubator for at least 36 h and a maximum of 48 h. **DO NOT** let the sample dry out. *Add a 100 mm petri dish filled with milliQ water to the bottom of the incubator for humidification.*

**13. Post Encoding Probe Hybridization Wash**
13.1. Remove the parafilm and add 5 mL Formamide Wash Buffer (PN 20300002).
13.2. Incubate at 47°C for 30 min in an incubator in a fume hood.
13.3. Aspirate the Formamide Wash Buffer. Add 5 mL Formamide Wash Buffer.
13.4. Incubate at 47°C for 30 min in an incubator in a fume hood.
At this point, the slides can be stored for up to 7 days in clearing premix at 37°C.

*OPTIONAL: Weighted gel embedding (to avoid tissue lifting during acquisition):*
*9.4. Clean a Gel Coverslip (PN 30200004) by spraying with RNaseZap solution and wiping with a Kimwipe, followed by spraying 70% ethanol and wiping with a Kimwipe.*
*9.5. Add 100 µL Gel Slick Solution onto the Gel Coverslip. Allow the Gel Slick Solution to evaporate for 10 min at room temperature. Wipe gently with a Kimwipe to remove any remaining film, liquid, or deposition from the glass. Use immediately after preparation.*
*9.6. Prepare Gel Embedding Solution:*

| Gel Embedding Solution | 1 sample |
|---|---|
| Gel Embedding Premix (PN 20300118) | 5 mL |
| 10% w/v ammonium persulfate solution | 25 µL |
| N,N,N′,N'-tetramethylethylenediamine | 2.5 µL |

*9.11. Aspirate the Sample Prep Wash Buffer. Retain 100 µL Gel Embedding Solution in a small tube. Add the remainder of the 5 mL Gel Embedding Solution to the sample, ensure the sample is fully covered, and incubate at room temperature for 1 min.*

*9.12. Using a pipette, transfer the majority of the Gel Embedding Solution to a waste tube (to monitor the gel formation).*

*9.13. Aspirate to dry the MERSCOPE FFPE Slide, leaving just enough liquid to cover the tissue section.*

*9.14. Add 50 µL of the retained Gel Embedding Solution on the tissue section.*

*9.15. Place the tips of one pair of tweezers on an area of the MERSCOPE FFPE Slide without touching the tissue section. Use tweezers to pick up the 20-mm Gel Slick-treated Gel Coverslip. With the Gel Slick-treated side facing down toward the tissue, place the edge of the Gel Coverslip against the tweezer tips resting on the MERSCOPE FFPE Slide, creating stability, and slowly lower the Gel Coverslip onto the tissue section to spread the Gel Embedding Solution. If needed, adjust the Gel Coverslip so it is positioned in the center of the MERSCOPE FFPE Slide. Gently press the Gel Coverslip to squeeze out excess Gel Embedding Solution and remove the extra Gel Embedding Solution by aspiration. Put a lid of 50 ml falcon with 2 ml RNAase-free water*

*9.16. Incubate at room temperature for 1.5 h.*

**14. Stain MERSCOPE slide with DAPI (right before imaging)**

14.1. Aspirate the Clearing Solution (from sample preparation) ensuring all Clearing Solution is aspirated from the petri dish.

14.2. Wash 2x with 5 mL Sample Prep Wash Buffer.

14.3. Gently shake the Verification Staining Reagent tube to ensure the reagent is well mixed and no precipitate is visible.

14.4. Add 3 mL Verification Staining Reagent, incubate 15 min on a rocker.

14.5. Wash 1x with 5 mL Formamide Wash Buffer, incubate 10 min.

14.6. Wash 1x with 5 mL Sample Prep Wash Buffer.

14.7. Proceed immediately to image the slide using the MERSCOPE Instrument (MERSCOPE Instrument User Guide 91600001)

#### D. Post-processing: segment the cells and generate expression matrixes

The original cell segmentation may not be satisfying, here, we provide a pre-trained cellpose2 model suitable for Xenopus embryo sections from gastrula to neurula stages. This model uses cell boundary staining 3 from the Vizgen cell segmentation kit as an input. As we have multiple samples in one section, we also offer a code for annotating the sample information.

The tutorial for the post-processing is on Github:
https://github.com/monsoro

Table S1. Summary table of cell type distribution by Model 1 and Model 2

| Model | unique_id | Stage | Endoderm_cells_count_only | Ectoderm_cells_count_only | Double_positive_count | Negative_count | Total_cells |
|---|---|---|---|---|---|---|---|
| **Model 1** | 1 | 9 | 31 | 195 | 48 | 47 | 321 |
| **Model 1** | 2 | 9 | 0 | 374 | 2 | 35 | 411 |
| **Model 1** | 3 | 9 | 9 | 42 | 2 | 193 | 246 |
| **Model 1** | 4 | 9 | 48 | 209 | 7 | 224 | 488 |
| **Model 1** | 5 | 9 | 0 | 182 | 0 | 49 | 231 |
| **Model 1** | 6 | 10.5 | 38 | 200 | 18 | 198 | 454 |
| **Model 1** | 7 | 10.5 | 79 | 107 | 4 | 171 | 361 |
| **Model 1** | 8 | 10.5 | 23 | 156 | 0 | 238 | 417 |
| **Model 1** | 9 | 10.5 | 37 | 279 | 0 | 393 | 709 |
| **Model 1** | 10 | 11 | 71 | 154 | 2 | 122 | 349 |
| **Model 1** | 11 | 11 | 36 | 161 | 8 | 164 | 369 |
| **Model 1** | 12 | 11 | 27 | 122 | 1 | 213 | 363 |
| **Model 1** | 13 | 11 | 125 | 328 | 3 | 506 | 962 |
| **Model 1** | 14 | 11 | 55 | 111 | 4 | 263 | 433 |
| **Model 1** | 15 | 12 | 63 | 280 | 7 | 453 | 803 |
| **Model 1** | 16 | 12 | 56 | 143 | 1 | 588 | 788 |
| **Model 1** | 17 | 12 | 29 | 70 | 1 | 314 | 414 |
| **Model 1** | 18 | 12 | 43 | 253 | 7 | 397 | 700 |
| **Model 1** | 19 | 12 | 16 | 90 | 2 | 351 | 459 |
| **Model 2** | 4 | 9 | 70 | 211 | 104 | 59 | 444 |
| **Model 2** | 5 | 9 | 3 | 536 | 9 | 54 | 602 |
| **Model 2** | 6 | 9 | 48 | 109 | 15 | 237 | 409 |
| **Model 2** | 7 | 9 | 134 | 251 | 53 | 199 | 637 |
| **Model 2** | 19 | 9 | 0 | 270 | 0 | 76 | 346 |
| **Model 2** | 8 | 10.5 | 166 | 138 | 19 | 130 | 453 |
| **Model 2** | 9 | 10.5 | 43 | 237 | 43 | 160 | 483 |
| **Model 2** | 10 | 10.5 | 107 | 194 | 5 | 457 | 763 |
| **Model 2** | 11 | 10.5 | 182 | 288 | 5 | 294 | 769 |
| **Model 2** | 14 | 11 | 274 | 212 | 15 | 293 | 794 |
| **Model 2** | 15 | 11 | 162 | 240 | 25 | 359 | 786 |
| **Model 2** | 16 | 11 | 114 | 186 | 1 | 442 | 743 |
| **Model 2** | 17 | 11 | 250 | 315 | 7 | 388 | 960 |
| **Model 2** | 18 | 11 | 123 | 192 | 21 | 312 | 648 |
| **Model 2** | 29 | 12 | 139 | 275 | 10 | 374 | 798 |
| **Model 2** | 30 | 12 | 167 | 166 | 1 | 603 | 937 |
| **Model 2** | 31 | 12 | 118 | 105 | 1 | 562 | 786 |
| **Model 2** | 32 | 12 | 103 | 244 | 11 | 425 | 783 |
| **Model 2** | 33 | 12 | 47 | 131 | 7 | 496 | 681 |

Table 2. Selected genes in the reference atlas and their expression across developmental stages

| Gene | Blastula st9 | Reference | Gastrula st10 | Reference | Gastrula st12 | Reference | Neurula | Reference | Tailbud | Reference | Xenbase link (expression page) |
|---|---|---|---|---|---|---|---|---|---|---|---|
| ACTC1.L | NA | | NA | | mesoderm<br>paraxial mesoderm<br>presomitic mesoderm | Della Gaspera B et al. (2012), https://www.xenbase.org/xenbase/literature/article.do?method=display&articleId=44993 | somite<br>presomitic mesoderm | Della Gaspera B et al. (2018), https://www.xenbase.org/xenbase/literature/article.do?method=display&articleId=55161 | somite<br>muscle<br>skeletal muscle<br>myotome (st22) | Meadows SM et al. (2008), Meadows SM et al. (2008) | https://www.xenbase.org/xenbase/gene/expression.do?method=displayGenePageExpression&geneId=480346&tabId=1 |
| CDH1.S | animal cap<br>presumptive<br>ectoderm<br>cell (st10) | Vacca B et al. (2018), https://www.xenbase.org/xenbase/literature/article.do?method=display&articleId=54471 | animal cap (from st10.5) | Nandadasa S et al. (2009), https://www.xenbase.org/xenbase/literature/article.do?method=display&articleId=39327 | ectoderm | Nandadasa S et al. (2009), https://www.xenbase.org/xenbase/literature/article.do?method=display&articleId=39327 | ectoderm | Levi G et al. (1991), https://www.xenbase.org/xenbase/literature/article.do?method=display&articleId=25171 | epidermis | Bharathan NK and Dickinson AJG (2019), https://www.xenbase.org/xenbase/literature/article.do?method=display&articleId=55821 | https://www.xenbase.org/xenbase/gene/expression.do?method=displayGenePageExpression&geneId=12531593&tabId=1 |
| CD44.L | NA | | NA | | NA | | somite | Ori M et al. (2006), https://www.xenbase.org/xenbase/literature/article.do?method=display&articleId=814 | somites, muscle progenitors | Ori M et al. (2006), https://www.xenbase.org/xenbase/literature/article.do?method=display&articleId=814 | https://www.xenbase.org/xenbase/gene/expression.do?method=displayGenePageExpression&geneId=487546&tabId=1 |
| CDX4.L | NA | | marginal zone | Nakamura Y et al. (2016),https://www.xenbase.org/xenbase/literature/article.do?method=display&articleId=52077 | circumblastoporal collar | Northrop JL and Kimelman D (1994), https://www.xenbase.org/xenbase/literature/article.do?method=display&articleId=21651 | posterior<br>mesoderm - st14 | Beck CW and Slack JM (1998), https://www.xenbase.org/xenbase/literature/article.do?method=display&articleId=15068 | posterior neural tube<br>posterior - st22 | Northrop JL and Kimelman D (1994), https://www.xenbase.org/xenbase/literature/article.do?method=display&articleId=21651 | https://www.xenbase.org/xenbase/gene/expression.do?method=displayGenePageExpression&geneId=482786&tabId=1 |
| CHRD.1.S | dorsal marginal zone | Harata A et al. (2019), https://www.xenbase.org/xenbase/literature/article.do?method=display&articleId=55162 | upper blastopore lip | Copyright © XDB3, Naoto Ueno, https://www.xenbase.org/xenbase/image/supplemental/XB-IMG-14267.jpg | mesoderm<br>dorsal marginal zone | Luque ME et al. (2008), https://www.xenbase.org/xenbase/literature/article.do?method=display&articleId=36961 | notochord - st18 | Chang C and Harland RM (2007), https://www.xenbase.org/xenbase/literature/article.do?method=display&articleId=36621 | notochord - st20 | de Almeida I et al. (2008), https://www.xenbase.org/xenbase/literature/article.do?method=display&articleId=37465 | https://www.xenbase.org/xenbase/gene/expression.do?method=displayGenePageExpression&geneId=480637&tabId=1 |
| DLX2.L | NA | | NA | | involuting marginal zone<br>involuted ventral mesoderm<br>circumblastoporal collar | Hufton AL et al. (2006), https://www.xenbase.org/xenbase/literature/article.do?method=display&articleId=256 | NA | | pharyngeal arch<br>cranial neural crest<br>mandibular crest<br>hyoid crest<br>anterior branchial crest<br>posterior branchial crest<br>migratory neural crest cell<br>periotic region<br>cranial placode<br>lens placode | Papalopulu N and Kintner C (1993), https://www.xenbase.org/xenbase/literature/article.do?method=display&articleId=22797 | https://www.xenbase.org/xenbase/gene/expression.do?method=displayGenePageExpression&geneId=852890&tabId=1 |
| DLX3.L | NA | | ectoderm<br>epidermis outer layer | Steiner AB et al. (2006), https://www.xenbase.org/xenbase/literature/article.do?method=display&articleId=34853 | ectoderm | Beanan MJ and Sargent TD (2000), https://www.xenbase.org/xenbase/literature/article.do?method=display&articleId=10631 | neural plate border<br>superficial<br>cement gland primordium<br>non-neural ectoderm | Schlosser G and Ahrens K (2004), https://www.xenbase.org/xenbase/literature/article.do?method=display&articleId=3353 | profundal placode<br>trigeminal placode<br>cement gland primordium<br>olfactory placode<br>otic placode - st21 - st22 | Schlosser G and Ahrens K (2004), https://www.xenbase.org/xenbase/literature/article.do?method=display&articleId=3353 | https://www.xenbase.org/xenbase/gene/expression.do?method=displayGenePageExpression&geneId=5879292&tabId=1 |
| EGR2.L | NA | | NA | | NA | | rhombomere R3<br>rhombomere R5<br>migratory neural crest cell - st14 | Read EM et al. (1998), https://www.xenbase.org/xenbase/literature/article.do?method=display&articleId=14332 | hindbrain<br>anterior branchial neural crest<br>neuroectoderm<br>rhombomere R3<br>rhombomere R5 | Dibner C et al. (2001), https://www.xenbase.org/xenbase/literature/article.do?method=display&articleId=8358 | https://www.xenbase.org/xenbase/gene/expression.do?method=displayGenePageExpression&geneId=853289&tabId=1 |
| ELAVL3.L | NA | | NA | | neuroectoderm<br>neural plate | Bang AG et al. (1999), https://www.xenbase.org/xenbase/literature/article.do?method=display&articleId=12578 | trigeminal placode<br>trunk neural crest<br>chordal neural plate<br>olfactory region<br>floor plate - st17 | Perron M et al. (1999), https://www.xenbase.org/xenbase/literature/article.do?method=display&articleId=12395 | neural tube<br>trigeminal placode<br>olfactory region - st20 | Perron M et al. (1999), https://www.xenbase.org/xenbase/literature/article.do?method=display&articleId=12395 | https://www.xenbase.org/xenbase/gene/expression.do?method=displayGenePageExpression&geneId=490863&tabId=1 |
| EPHA2.L | NA | | NA | | NA | | hindbrain<br>rhombomere R4 - st18 | Elkouby YM et al. (2012), https://www.xenbase.org/xenbase/literature/article.do?method=display&articleId=44950 | anterior branchial crest - st20 | Smith A et al. (1997), https://www.xenbase.org/xenbase/literature/article.do?method=display&articleId=16188 | https://www.xenbase.org/xenbase/gene/expression.do?method=displayGenePageExpression&geneId=5761258&tabId=1 |
| EPHA4.L | NA | | mesoderm<br>involuted dorsal mesoderm<br>involuted ventral mesoderm | Park EC et al. (2011), https://www.xenbase.org/xenbase/literature/article.do?method=display&articleId=42478 | NA | | brain<br>neural tube<br>neuroectoderm<br>eye primordium<br>preplacodal ectoderm<br>presumptive rhombomere<br>rhombomere R3<br>rhombomere R5 - st15 | Maldonado-Agurto R et al. (2011), https://www.xenbase.org/xenbase/literature/article.do?method=display&articleId=43476 | branchial crest<br>hyoid crest<br>eye<br>head<br>hindbrain<br>cranial neural crest - st21 to st24 | Smith A et al. (1997), https://www.xenbase.org/xenbase/literature/article.do?method=display&articleId=16188 | https://www.xenbase.org/xenbase/gene/expression.do?method=displayGenePageExpression&geneId=1015802&tabId=1 |
| EYA1.L | animal cap<br>ectoderm | Neilson KM et al. (2010), https://www.xenbase.org/xenbase/literature/article.do?method=display&articleId=42372 | animal cap<br>ectoderm | Neilson KM et al. (2010), https://www.xenbase.org/xenbase/literature/article.do?method=display&articleId=42372 | animal cap<br>ectoderm | Neilson KM et al. (2010), https://www.xenbase.org/xenbase/literature/article.do?method=display&articleId=42372 | anterior neural fold<br>trigeminal placode<br>neural plate - st 13- st14 | Schlosser G and Ahrens K (2004), https://www.xenbase.org/xenbase/literature/article.do?method=display&articleId=3353 | trigeminal placode<br>olfactory placode<br>otic placode | Schlosser G and Ahrens K (2004), https://www.xenbase.org/xenbase/literature/article.do?method=display&articleId=3353 | https://www.xenbase.org/xenbase/gene/expression.do?method=displayGenePageExpression&geneId=479278&tabId=1 |
| EYA2.S | NA | | NA | | NA | | neural plate<br>preplacodal ectoderm -st17 | Neilson KM et al. (2010), https://www.xenbase.org/xenbase/literature/article.do?method=display&articleId=42372 | preplacodal ectoderm<br>trigeminal placode<br>adenohypophyseal placode<br>olfactory placode<br>neurogenic placode<br>cranial placode - st20 | Neilson KM et al. (2010), https://www.xenbase.org/xenbase/literature/article.do?method=display&articleId=42372 | https://www.xenbase.org/xenbase/gene/expression.do?method=displayGenePageExpression&geneId=487843&tabId=1 |
| FOXC2.L | NA | | NA | | presumptive paraxial mesoderm | El-Hodiri H et al. (2001), https://www.xenbase.org/xenbase/literature/article.do?method=display&articleId=9235 | paraxial mesoderm | El-Hodiri H et al. (2001), https://www.xenbase.org/xenbase/literature/article.do?method=display&articleId=9235 | mesoderm<br>head<br>somite<br>sclerotome<br>presomitic mesoderm<br>head mesenchyme (st 22) | Soeren S Lienkamp , https://www.xenbase.org/xenbase/image/supplemental/XB-IMG-156303.jpg | https://www.xenbase.org/xenbase/gene/expression.do?method=displayGenePageExpression&geneId=481381&tabId=1 |
| GATA2.L | animal cap | Pieper M et al. (2012), https://www.xenbase.org/xenbase/literature/article.do?method=display&articleId=44816 | ectoderm<br>animal cap | Read EM et al. (1998), https://www.xenbase.org/xenbase/literature/article.do?method=display&articleId=14332 | non-neural ectoderm<br>epidermis | Pieper M et al. (2012), https://www.xenbase.org/xenbase/literature/article.do?method=display&articleId=44816 | epidermis<br>ventral<br>ectoderm -st15 | Bellefroid EJ et al. (1998), https://www.xenbase.org/xenbase/literature/article.do?method=display&articleId=15410 | ventral blood island<br>cardinal vein<br>periocular region<br>head vasculature - st23 | Liu F et al. (2008), https://www.xenbase.org/xenbase/literature/article.do?method=display&articleId=38293 | https://www.xenbase.org/xenbase/gene/expression.do?method=displayGenePageExpression&geneId=478164&tabId=1 |
| GBX2.2.L | NA | | NA | | posterior neuroectoderm | Li B et al. (2009), https://www.xenbase.org/xenbase/literature/article.do?method=display&articleId=40487 | midbrain-hindbrain boundary<br>neural crest<br>epidermis | Glavic A et al. (2002), https://www.xenbase.org/xenbase/literature/article.do?method=display&articleId=7392 | midbrain-hindbrain boundary<br>rhombomere<br>head endoderm<br>foregut<br>eye<br>cranial neural crest | von Bubnoff A et al. (1996), https://www.xenbase.org/xenbase/literature/article.do?method=display&articleId=18598 | https://www.xenbase.org/xenbase/gene/expression.do?method=displayGenePageExpression&geneId=852623&tabId=1 |

| | | | | | | | | | | | |
|---|---|---|---|---|---|---|---|---|---|---|---|
| HNF1B.L | NA | | endoderm<br>upper blastopore lip | Gere-Becker MB et al. (2018)<br>https://www.xenbase.org/xenbase/literature/article.do?method=display&articleId=54918 | endoderm<br>upper blastopore lip | Gere-Becker MB et al. (2018)<br>https://www.xenbase.org/xenbase/literature/article.do?method=display&articleId=54918 | lateral plate mesoderm<br>dorsal lateral plate mesoderm | Buisson I et al. (2015),<br>https://www.xenbase.org/xenbase/literature/article.do?method=display&articleId=49971 | pronephric mesenchyme | Marracci S et al. (2016),<br>https://www.xenbase.org/xenbase/literature/article.do?method=display&articleId=51918 | https://www.xenbase.org/xenbase/gene/expression.do?method=displayGenePageExpression&geneId=485651&tabId=1 |
| HOXA3.S | NA | | NA | | NA | | presumptive rhombomere | Bae CJ et al. (2015),<br>https://www.xenbase.org/xenbase/literature/article.do?method=display&articleId=50095 | hindbrain<br>cranial neural crest<br>spinal cord<br>anterior branchial crest<br>otic vesicle | McNulty CL et al. (2005),<br>https://www.xenbase.org/xenbase/literature/article.do?method=display&articleId=1812 | https://www.xenbase.org/xenbase/gene/expression.do?method=displayGenePageExpression&geneId=482266&tabId=1 |
| HOXB6.S | NA | | NA | | NA | | presumptive rhombomere | https://www.xenbase.org/xenbase/community/viewPerson.do?method=display&personId=3172 | tail bud<br>eye<br>pronephric kidney<br>head<br>spinal cord<br>pharyngeal region<br>intermediate mesoderm - st28 | https://www.xenbase.org/xenbase/community/viewPerson.do?method=display&personId=3172 | https://www.xenbase.org/xenbase/gene/expression.do?method=displayGenePageExpression&geneId=1006004&tabId=1 |
| HOXB9.S | NA | | NA | | circumblastoporal collar | Zhu K et al. (2017),<br>https://www.xenbase.org/xenbase/literature/article.do?method=display&articleId=53181 | neural tube<br>posterior | Chang C and Harland RM (2007),<br>https://www.xenbase.org/xenbase/literature/article.do?method=display&articleId=36621 | neural tube<br>posterior | Dibner C et al. (2001),<br>https://www.xenbase.org/xenbase/literature/article.do?method=display&articleId=8358 | https://www.xenbase.org/xenbase/gene/expression.do?method=displayGenePageExpression&geneId=963093&tabId=1 |
| HOXC10.L | NA | | NA | | NA | | presomitic mesoderm<br>caudal presomitic mesoderm - st18 | Young JJ et al. (2014),<br>https://www.xenbase.org/xenbase/literature/article.do?method=display&articleId=48813 | posterior<br>presomitic mesoderm<br>somite<br>trunk somite<br>tail somite<br>tail bud<br>posterior neural tube | Christen B et al. (2003)<br>https://www.xenbase.org/xenbase/literature/article.do?method=display&articleId=5825, | https://www.xenbase.org/xenbase/gene/expression.do?method=displayGenePageExpression&geneId=486914&tabId=1 |
| HOXC9.S | NA | | NA | | NA | | NA | | NA | | https://www.xenbase.org/xenbase/gene/expression.do?method=displayGenePageExpression&geneId=482329&tabId=1 |
| HOXD11.L | NA | | NA | | NA | | NA | | tail<br>tail tip<br>chordoneural hinge<br>posterior<br>spinal cord<br>blood vessel<br>intersomitic vessel -st33 | https://www.xenbase.org/xenbase/community/viewPerson.do?method=display&personId=3696 | https://www.xenbase.org/xenbase/gene/expression.do?method=displayGenePageExpression&geneId=485843&tabId=1 |
| HOXD3.L | NA | | NA | | circumblastoporal collar | https://www.xenbase.org/xenbase/community/lab.do?method=display&labId=241 | hindbrain<br>mesoderm<br>neuroectoderm | Lloret-Vilaspasa F et al. (2010),<br>https://www.xenbase.org/xenbase/literature/article.do?method=display&articleId=41268 | hindbrain<br>rhombomere R5<br>neural tube<br>neuroectoderm | Lloret-Vilaspasa F et al. (2010),<br>https://www.xenbase.org/xenbase/literature/article.do?method=display&articleId=41268 | https://www.xenbase.org/xenbase/gene/expression.do?method=displayGenePageExpression&geneId=478461&tabId=1 |
| KRT12.4.L | NA | | ectoderm<br>animal<br>hemisphere | Copyright © XDB3, Naoto Ueno<br>https://www.xenbase.org/xenbase/ViewImageActionNonAdmin.do?imageId=13582 | ectoderm<br>epidermis | Li B et al. (2009),<br>https://www.xenbase.org/xenbase/literature/article.do?method=display&articleId=40487 | epidermis -st18 | Whittington N et al. (2015),<br>https://www.xenbase.org/xenbase/literature/article.do?method=display&articleId=49962 | non-neural ectoderm<br>epidermis -st22 | Pieper M et al. (2012),<br>https://www.xenbase.org/xenbase/literature/article.do?method=display&articleId=44816 | https://www.xenbase.org/xenbase/gene/expression.do?method=displayGenePageExpression&geneId=6539689&tabId=1 |
| MAFB.S | NA | | NA | | NA | | presumptive rhombomere | Chung HA et al. (2014),<br>https://www.xenbase.org/xenbase/literature/article.do?method=display&articleId=48815 | somite<br>ventral blood island dorsal<br>neural tube neural crest | Ishibashi S and Yasuda K (2001),<br>https://www.xenbase.org/xenbase/literature/article.do?method=display&articleId=9470 | https://www.xenbase.org/xenbase/gene/expression.do?method=displayGenePageExpression&geneId=6085858&tabId=1 |
| MSX1.L | NA | | ventro-lateral<br>marginal zone | Freeman SD et al. (2008),<br>https://www.xenbase.org/xenbase/literature/article.do?method=display&articleId=38063 | neural crest | Li B et al. (2009),<br>https://www.xenbase.org/xenbase/literature/article.do?method=display&articleId=40487 | ectoderm<br>neural plate border<br>pre-chordal neural plate border<br>chordal neural plate border<br>preplacodal ectoderm -st16 | Dichmann DS and Harland RM (2011),<br>https://www.xenbase.org/xenbase/literature/article.do?method=display&articleId=42231 | neuroectoderm<br>neural crest<br>neural plate - st21 | Glavic A et al. (2004),<br>https://www.xenbase.org/xenbase/literature/article.do?method=display&articleId=4249 | https://www.xenbase.org/xenbase/gene/expression.do?method=displayGenePageExpression&geneId=490377&tabId=1 |
| MSX2.L | NA | | ventral marginal<br>zone | Popov IK et al. (2017),<br>https://www.xenbase.org/xenbase/literature/article.do?method=display&articleId=52133 | neuroectoderm<br>neural plate border | Khadka D et al. (2006),<br>https://www.xenbase.org/xenbase/literature/article.do?method=display&articleId=495 | neural crest<br>non-neural ectoderm | Khadka D et al. (2006),<br>https://www.xenbase.org/xenbase/literature/article.do?method=display&articleId=495 | head<br>mandibular arch<br>neural crest -st28 | Kennedy AE and Dickinson AJ (2012),<br>https://www.xenbase.org/xenbase/literature/article.do?method=display&articleId=44961 | https://www.xenbase.org/xenbase/gene/expression.do?method=displayGenePageExpression&geneId=852973&tabId=1 |
| MYL1.L | NA | | NA | | NA | | NA | | somite | Neuner R et al. (2009),<br>https://www.xenbase.org/xenbase/literature/article.do?method=display&articleId=38602 | https://www.xenbase.org/xenbase/gene/expression.do?method=displayGenePageExpression&geneId=970332&tabId=1 |
| MYOD1.S | NA | | mesoderm<br>dorso-lateral<br>marginal zone<br>(st10.5) | Copyright © CNRS UMR 8080, Nicolas Pollet,<br>https://www.xenbase.org/xenbase/ViewImageActionNonAdmin.do?imageId=29373 | presumptive<br>paraxial mesoderm | Sempou E et al. (2018),<br>https://www.xenbase.org/xenbase/literature/article.do?method=display&articleId=55550 | presomitic mesoderm<br>paraxial mesoderm (st18) | Chang C and Harland RM (2007),<br>https://www.xenbase.org/xenbase/literature/article.do?method=display&articleId=36621 | presomitic mesoderm | Della Gaspera B et al. (2012),<br>https://www.xenbase.org/xenbase/literature/article.do?method=display&articleId=44993 | https://www.xenbase.org/xenbase/gene/expression.do?method=displayGenePageExpression&geneId=1017497&tabId=1 |
| NOG.L | dorsal marginal<br>zone | Wagner G et al. (2017),<br>https://www.xenbase.org/xenbase/literature/article.do?method=display&articleId=53676 | dorsal marginal<br>zone<br>upper blastopore<br>lip | Fletcher RB and Harland RM (2008),<br>https://www.xenbase.org/xenbase/literature/article.do?method=display&articleId=37484 | axial mesoderm | Oelgeschläger M et al. (2003),<br>https://www.xenbase.org/xenbase/literature/article.do?method=display&articleId=5765 | axial mesoderm<br>notochord (st18) | Fletcher RB and Harland RM (2008),<br>https://www.xenbase.org/xenbase/literature/article.do?method=display&articleId=37484 | forebrain<br>notochord<br>branchial crest- st 21 | Fletcher RB et al. (2004),<br>https://www.xenbase.org/xenbase/literature/article.do?method=display&articleId=2676 | https://www.xenbase.org/xenbase/gene/expression.do?method=displayGenePageExpression&geneId=487723&tabId=1 |
| OLIG3.S | NA | | NA | | NA | | NA | | NA | | https://www.xenbase.org/xenbase/gene/expression.do?method=displayGenePageExpression&geneId=919788&tabId=1 |
| OLIG4.L | NA | | NA | | NA | | paraxial mesoderm<br>intermediate mesoderm<br>chordal neural plate | Martynova NY et al. (2013),<br>https://www.xenbase.org/xenbase/literature/article.do?method=display&articleId=47095 | chordal neural plate<br>roof plate | https://www.xenbase.org/xenbase/community/lab.do?method=display&labId=23 | https://www.xenbase.org/xenbase/gene/expression.do?method=displayGenePageExpression&geneId=484735&tabId=1 |
| OTX2.S | NA | | upper blastopore<br>lip<br>dorsal | Fletcher RB and Harland RM (2008),<br>https://www.xenbase.org/xenbase/literature/article.do?method=display&articleId=37484 | neural plate | Li B et al. (2009),<br>https://www.xenbase.org/xenbase/literature/article.do?method=display&articleId=40487 | neuroectoderm<br>diencephalon<br>telencephalon<br>pre-chordal neural plate<br>anterior neural fold | Read EM et al. (1998),<br>https://www.xenbase.org/xenbase/literature/article.do?method=display&articleId=14332 | forebrain<br>optic vesicle | Vignali R et al. (2000),<br>https://www.xenbase.org/xenbase/literature/article.do?method=display&articleId=10513 | https://www.xenbase.org/xenbase/gene/expression.do?method=displayGenePageExpression&geneId=485219&tabId=1 |
| PAX3.S | NA | | NA | | neural plate border | Hatch VL et al. (2016),<br>https://www.xenbase.org/xenbase/literature/article.do?method=display&articleId=52355 | neural plate<br>pre-chordal neural plate<br>chordal neural plate<br>profundal placode -st16 | Schlosser G and Ahrens K (2004),<br>https://www.xenbase.org/xenbase/literature/article.do?method=display&articleId=3353 | neural plate<br>neural tube<br>posterior neural tube<br>anterior neural tube<br>hatching gland -st18 to st21 | Borchers A et al. (2006),<br>https://www.xenbase.org/xenbase/literature/article.do?method=display&articleId=521 | https://www.xenbase.org/xenbase/gene/expression.do?method=displayGenePageExpression&geneId=482739&tabId=1 |
| PAX6.S | NA | | NA | | optic field<br>prechordal plate | Wei S et al. (2012),<br>https://www.xenbase.org/xenbase/literature/article.do?method=display&articleId=44671 | neuroectoderm<br>eye primordium<br>neural plate<br>chordal neural plate | Mamada H et al. (2009),<br>https://www.xenbase.org/xenbase/literature/article.do?method=display&articleId=39139 | optic vesicle<br>anterior neural tube | Escalante-Alcalde D et al. (2003),<br>https://www.xenbase.org/xenbase/literature/article.do?method=display&articleId=4825 | https://www.xenbase.org/xenbase/gene/expression.do?method=displayGenePageExpression&geneId=484087&tabId=1 |

| | | | | | | | | | | | |
|---|---|---|---|---|---|---|---|---|---|---|---|
| PAX7.L | NA | | NA | | NA | | neural plate<br>paraxial mesoderm | Maczkowiak F et al. (2010), https://www.xenbase.org/xenbase/literature/article.do?method=display&articleId=41089 | roof plate<br>midbrain<br>hindbrain<br>myotome<br>paraxial mesoderm | Maczkowiak F et al. (2010), https://www.xenbase.org/xenbase/literature/article.do?method=display&articleId=41089 | https://www.xenbase.org/xenbase/gene/expression.do?method=displayGenePageExpression&geneId=487139&tabId=1 |
| PITX1.L | NA | | NA | | NA | | ectoderm<br>cement gland primordium | Chang W et al. (2001), https://www.xenbase.org/xenbase/literature/article.do?method=display&articleId=9612 | anterior neural tube<br>cement gland<br>eye primordium<br>stomodeal-hypophyseal primordium | Chang W et al. (2001), https://www.xenbase.org/xenbase/literature/article.do?method=display&articleId=9612 | https://www.xenbase.org/xenbase/gene/expression.do?method=displayGenePageExpression&geneId=485440&tabId=1 |
| RAX.L | NA | | NA | | neuroectoderm<br>neural plate<br>sensorial layer | Andreazzoli M et al. (2003), https://www.xenbase.org/xenbase/literature/article.do?method=display&articleId=4688 | neural plate<br>anterior<br>eye primordium | Zuber ME et al. (2003), https://www.xenbase.org/xenbase/literature/article.do?method=display&articleId=4775 | optic vesicle<br>retina | Lee HX et al. (2006), https://www.xenbase.org/xenbase/literature/article.do?method=display&articleId=841 | https://www.xenbase.org/xenbase/gene/expression.do?method=displayGenePageExpression&geneId=492664&tabId=1 |
| SIX1.L | NA | | endomesoderm<br>dorsal endoderm<br>dorsal<br>endomesoderm | Mukherjee S et al. (2020), https://www.xenbase.org/xenbase/literature/article.do?method=display&articleId=57340 | pre-chordal neural plate<br>anterior neural ridge | Ahrens K and Schlosser G (2005), https://www.xenbase.org/xenbase/literature/article.do?method=display&articleId=1139 | trigeminal placode<br>anterior neural fold<br>profundal placode | Schlosser G and Ahrens K (2004), https://www.xenbase.org/xenbase/literature/article.do?method=display&articleId=3353 | trigeminal placode<br>olfactory placode<br>otic placode -st21/22 | Schlosser G and Ahrens K (2004), https://www.xenbase.org/xenbase/literature/article.do?method=display&articleId=3353 | https://www.xenbase.org/xenbase/gene/expression.do?method=displayGenePageExpression&geneId=480716&tabId=1 |
| SNAI1.S | NA | | marginal zone<br>mesoderm | https://www.xenbase.org/xenbase/community/viewPerson.do?method=display&personId=752 | neural plate border<br>presumptive paraxial mesoderm | O'Donnell M et al. (2006), https://www.xenbase.org/xenbase/literature/article.do?method=display&articleId=34474 | cranial neural crest | Lee YH et al. (2004), https://www.xenbase.org/xenbase/literature/article.do?method=display&articleId=2920 | cranial neural crest<br>migratory neural crest cell<br>neural plate border<br>pre-chordal neural plate border<br>chordal neural plate border -st21 | Peres JN and Durston AJ (2006), https://www.xenbase.org/xenbase/literature/article.do?method=display&articleId=870 | https://www.xenbase.org/xenbase/gene/expression.do?method=displayGenePageExpression&geneId=479969&tabId=1 |
| SNAI2.L | NA | | NA | | neural plate border | Li B et al. (2009), https://www.xenbase.org/xenbase/literature/article.do?method=display&articleId=40487 | neural crest | Glavic A et al. (2004), https://www.xenbase.org/xenbase/literature/article.do?method=display&articleId=4249 | neural crest | LaBonne C and Bronner-Fraser M (2000), https://www.xenbase.org/xenbase/literature/article.do?method=display&articleId=11164 | https://www.xenbase.org/xenbase/gene/expression.do?method=displayGenePageExpression&geneId=487370&tabId=1 |
| SOX10.L | NA | | NA | | NA | | cranial neural crest | Aoki Y et al. (2003), https://www.xenbase.org/xenbase/literature/article.do?method=display&articleId=5136 | neural crest | Kumasaka M et al. (2005), https://www.xenbase.org/xenbase/literature/article.do?method=display&articleId=1624 | https://www.xenbase.org/xenbase/gene/expression.do?method=displayGenePageExpression&geneId=480303&tabId=1 |
| SOX17A.S | vegetal<br>hemisphere<br>endoderm | Zhang C et al. (2005), https://www.xenbase.org/xenbase/literature/article.do?method=display&articleId=2400 | endoderm | Copyright © Zorn Lab, Aaron M Zorn, https://www.xenbase.org/xenbase/ViewImageActionNonAdmin.do?imageId=38628 | endoderm | Lim JW et al. (2011), https://www.xenbase.org/xenbase/ViewImageActionNonAdmin.do?imageId=49619 | endoderm | Copyright © Zorn Lab, Aaron M Zorn https://www.xenbase.org/xenbase/ViewImageActionNonAdmin.do?imageId=38630 | primary heart field<br>endoderm st 23 | Ito Y et al. (2008), https://www.xenbase.org/xenbase/literature/article.do?method=display&articleId=37722 | https://www.xenbase.org/xenbase/gene/expression.do?method=displayGenePageExpression&geneId=484294&tabId=1 |
| SOX2.L | animal pole | Sempou E et al. (2022), https://www.xenbase.org/xenbase/literature/article.do?method=display&articleId=59330 | neuroectoderm | Gao Y et al. (2015), https://www.xenbase.org/xenbase/literature/article.do?method=display&articleId=51023 | neural plate<br>neuroectoderm | Agüero TH et al. (2012), https://www.xenbase.org/xenbase/literature/article.do?method=display&articleId=44823 | neural plate<br>pre-chordal neural plate<br>chordal neural plate (st 18) | Chang C and Harland RM (2007), https://www.xenbase.org/xenbase/literature/article.do?method=display&articleId=36621 | optic vesicle<br>brain<br>forebrain<br>midbrain<br>hindbrain<br>olfactory region<br>spinal cord<br>trigeminal ganglion (st 22) | Wills AE et al. (2010), https://www.xenbase.org/xenbase/literature/article.do?method=display&articleId=40657 | https://www.xenbase.org/xenbase/gene/expression.do?method=displayGenePageExpression&geneId=484552&tabId=1 |
| SOX3.S | animal<br>hemisphere | Zhang C et al. (2005), https://www.xenbase.org/xenbase/literature/article.do?method=display&articleId=2400 | neuroectoderm<br>dorsal | Li HY et al. (2006), https://www.xenbase.org/xenbase/literature/article.do?method=display&articleId=954 | ectoderm<br>neural plate | Ahrens K and Schlosser G (2005), https://www.xenbase.org/xenbase/literature/article.do?method=display&articleId=1139 | neural plate<br>pre-chordal neural plate<br>chordal neural plate<br>trigeminal placode (st18) | Chang C and Harland RM (2007), https://www.xenbase.org/xenbase/literature/article.do?method=display&articleId=36621 | otic vesicle<br>olfactory placode<br>optic vesicle<br>profundus ganglion<br>trigeminal ganglion<br>forebrain<br>midbrain<br>hindbrain (st 22) | Schlosser G and Ahrens K (2004), https://www.xenbase.org/xenbase/literature/article.do?method=display&articleId=3353 | https://www.xenbase.org/xenbase/gene/expression.do?method=displayGenePageExpression&geneId=484814&tabId=1 |
| SOX8.L | NA | | NA | | ectoderm<br>neuroectoderm<br>neural crest | | Neural crest | | Neural crest | | https://www.xenbase.org/xenbase/gene/expression.do?method=displayGenePageExpression&geneId=480864&tabId=1 |
| SOX9.S | NA | | NA | | Neural crest | | Neural crest | | Neural crest | | https://www.xenbase.org/xenbase/gene/expression.do?method=displayGenePageExpression&geneId=1034768&tabId=1 |
| TFAP2A.L | animal<br>hemisphere | | ectoderm | | non-neural ectoderm | | neural crest<br>cranial neural crest (st18) | | neural crest<br>central nervous system (st22) | | https://www.xenbase.org/xenbase/gene/expression.do?method=displayGenePageExpression&geneId=481200&tabId=1 |
| TFAP2C.L | NA | | NA | | NA | | epidermis<br>pronephric mesenchyme<br>neural crest | Zhang Y et al. (2006),https://www.xenbase.org/xenbase/literature/article.do?m\ethod=display&articleId=837 | epidermis<br>pronephric mesenchyme<br>neural crest | Zhang Y et al. (2006),https://www.xenbase.org/xenbase/literature/article.do?m\ethod=display&articleId=837 | https://www.xenbase.org/xenbase/gene/expression.do?method=displayGenePageExpression&geneId=488670&tabId=1 |
| TP63.L | NA | | NA | | NA | | epidermis<br>dorsal<br>cement gland primordium | Barton CE et al. (2009), https://www.xenbase.org/xenbase/literature/article.do?method=display&articleId=39341 | epidermis inner layer | Cibois M et al. (2015), https://www.xenbase.org/xenbase/literature/article.do?method=display&articleId=50852 | https://www.xenbase.org/xenbase/gene/expression.do?method=displayGenePageExpression&geneId=479867&tabId=1 |
| VENTX2.1.L | NA | | ventral marginal zone<br>ventro-lateral marginal zone | Bhushan, et al. 1998, https://www.xenbase.org/xenbase/literature/article.do?method=display&articleId=14441 | ectoderm<br>neuroectoderm<br>involuted ventral mesoderm | Schmidt, et al. 1996, https://www.xenbase.org/xenbase/literature/article.do?method=display&articleId=18148 | mesoderm<br>eye primordium<br>cement gland primordium<br>posterior endoderm<br>posterior neural tube<br>neural tube<br>endomesoderm st18-19 | Copyright © Harland Lab https://www.xenbase.org/xenbase/ViewImageActionNonAdmin.do?imageId=37734 | posterior wall of neurenteric canal<br>eye<br>tail bud<br>pharyngeal region - st29/30 | Gawantka V et al. (1998), https://www.xenbase.org/xenbase/literature/article.do?method=display&articleId=13902 | https://www.xenbase.org/xenbase/gene/expression.do?method=displayGenePageExpression&geneId=919663&tabId=1 |
| ZIC1.S | NA | | neuroectoderm<br>inner layer<br>neuroectoderm<br>outer layer (st 10.25) | Steiner, et al. 2006 https://www.xenbase.org/xenbase/literature/article.do?method=display&articleId=34853 | neuroectoderm | Hatch, et al. 2016, https://www.xenbase.org/xenbase/literature/article.do?method=display&articleId=52355 | neural plate border<br>cranial neural crest (st18) | Pieper, et al. 2012 https://www.xenbase.org/xenbase/literature/article.do?method=display&articleId=44816 | somite<br>forebrain<br>brain<br>midbrain<br>hindbrain (st 22) | Almeida AD et al. (2010), Almeida AD et al. (2010) | https://www.xenbase.org/xenbase/gene/expression.do?method=displayGenePageExpression&geneId=481981&tabId=1 |
| TBXT.S | dorsal marginal<br>zone | Anderson GA et al. (2017), https://www.xenbase.org/xenbase/literature/article.do?method=display&articleId=54096 | marginal zone | Cornesse Y et al. (2005), https://www.xenbase.org/xenbase/literature/article.do?method=display&articleId=2541 | marginal zone | Northrop J et al. (1995), https://www.xenbase.org/xenbase/literature/article.do?method=display&articleId=19096 | circumblastoporal collar<br>mesenchyme<br>posterior<br>axial mesoderm -st14 | Beck CW and Slack JM (1998), https://www.xenbase.org/xenbase/literature/article.do?method=display&articleId=15068 | tail bud<br>chordoneural hinge<br>posterior wall of neurenteric canal - st29/30 | Beck CW and Slack JM (1998), https://www.xenbase.org/xenbase/literature/article.do?method=display&articleId=15068 | https://www.xenbase.org/xenbase/gene/expression.do?method=displayGenePageExpression&geneId=478788&tabId=1 |

Table 3. Evaluation of the gene panel efficiency

| | Gastrula (Stage 12) | | | | | | | Tailbud (Stage 22) | | | | | | | |
|---|---|---|---|---|---|---|---|---|---|---|---|---|---|---|---|
| gene | total_counts | mean_counts | number of cells with gene expression | percentage % of cells without gene expression | percentage of single transcript event (%) (Potential background) | number of single transcript event | Moran's I score | total_counts | mean_counts | number of cells with gene expression | percentage % of cells without gene expression) | percentage of single transcript event (%) (Potential background) | number of single transcript event | Moran's I score | Efficiency |
| ACTC1.L | 3211 | 0,806 | 944 | 76,311 | 14,329 | 571 | 0,538 | 287277 | 43,096 | 3350 | 49,745 | 19,442 | 1296 | 0,925 | High |
| CD44.L | 811 | 0,204 | 690 | 82,685 | 14,705 | 586 | 0,064 | 8146 | 1,222 | 2294 | 65,587 | 18,062 | 1204 | 0,680 | High |
| CDH1.S | 4405 | 1,105 | 1582 | 60,301 | 18,821 | 750 | 0,384 | 62357 | 9,354 | 4113 | 38,299 | 16,067 | 1071 | 0,722 | High |
| CDX4.L | 14883 | 3,735 | 3238 | 18,745 | 18,068 | 720 | 0,394 | 1950 | 0,293 | 990 | 85,149 | 11,071 | 738 | 0,632 | High |
| CHRD.1.S | 28178 | 7,071 | 1420 | 64,366 | 13,099 | 522 | 0,858 | 29376 | 4,407 | 3311 | 50,330 | 16,382 | 1092 | 0,783 | High |
| DLX2.L | 357 | 0,090 | 284 | 92,873 | 5,847 | 233 | 0,106 | 7826 | 1,174 | 1201 | 81,983 | 8,191 | 546 | 0,741 | High |
| DLX3.L | 2761 | 0,693 | 1519 | 61,882 | 22,836 | 910 | 0,229 | 16834 | 2,525 | 4173 | 37,399 | 24,107 | 1607 | 0,593 | High |
| EGR2.L | 1176 | 0,295 | 909 | 77,189 | 17,591 | 701 | 0,063 | 4525 | 0,679 | 1522 | 77,168 | 16,982 | 1132 | 0,592 | HIGH |
| ELAVL3.L | 4133 | 1,037 | 1365 | 65,747 | 16,612 | 662 | 0,520 | 11446 | 1,717 | 1460 | 78,098 | 12,541 | 836 | 0,684 | HIGH |
| EPHA2.L | 36537 | 9,169 | 3191 | 19,925 | 14,203 | 566 | 0,694 | 42850 | 6,428 | 4778 | 28,323 | 20,942 | 1396 | 0,672 | HIGH |
| EPHA4.L | 31999 | 8,030 | 3442 | 13,626 | 12,622 | 503 | 0,542 | 48285 | 7,243 | 4687 | 29,688 | 16,982 | 1132 | 0,672 | HIGH |
| EYA1.L | 656 | 0,165 | 517 | 87,026 | 10,740 | 428 | 0,096 | 9414 | 1,412 | 3287 | 50,690 | 19,892 | 1326 | 0,377 | HIGH |
| EYA2.S | 1139 | 0,286 | 863 | 78,344 | 16,612 | 662 | 0,095 | 23035 | 3,456 | 3957 | 40,639 | 15,887 | 1059 | 0,613 | HIGH |
| FOXC2.L | 10487 | 2,632 | 2019 | 49,335 | 21,054 | 839 | 0,578 | 33093 | 4,964 | 3741 | 43,879 | 16,007 | 1067 | 0,663 | HIGH |
| GATA2.L | 18647 | 4,679 | 2225 | 44,166 | 16,512 | 658 | 0,664 | 45952 | 6,893 | 4568 | 31,473 | 14,446 | 963 | 0,656 | HIGH |
| GBX2.2.L | 1194 | 0,300 | 744 | 81,330 | 12,622 | 503 | 0,324 | 5034 | 0,755 | 2158 | 67,627 | 17,402 | 1160 | 0,386 | HIGH |
| HNF1B.L | 729 | 0,183 | 541 | 86,424 | 11,041 | 440 | 0,227 | 9847 | 1,477 | 2762 | 58,566 | 19,082 | 1272 | 0,560 | HIGH |
| HOXA3.S | 1205 | 0,302 | 927 | 76,738 | 18,018 | 718 | 0,037 | 11564 | 1,735 | 3958 | 40,624 | 23,552 | 1570 | 0,391 | HIGH |
| HOXB6.S | 4686 | 1,176 | 1970 | 50,565 | 22,258 | 887 | 0,282 | 4199 | 0,630 | 1510 | 77,348 | 14,146 | 943 | 0,560 | HIGH |
| HOXB9.S | 696 | 0,175 | 478 | 88,005 | 8,683 | 346 | 0,211 | 4684 | 0,703 | 932 | 86,019 | 8,086 | 539 | 0,786 | HIGH |
| HOXC10.L | 994 | 0,249 | 774 | 80,577 | 15,182 | 605 | 0,087 | 1721 | 0,258 | 1326 | 80,108 | 15,692 | 1046 | 0,138 | MEDIUM |
| HOXC9.S | 1442 | 0,362 | 907 | 77,240 | 15,583 | 621 | 0,142 | 909 | 0,136 | 731 | 89,034 | 9,091 | 606 | 0,037 | MEDIUM |
| HOXD11.L | 428 | 0,107 | 376 | 90,565 | 8,331 | 332 | 0,108 | 1407 | 0,211 | 1192 | 82,118 | 15,287 | 1019 | 0,033 | MEDIUM |
| HOXD3.L | 972 | 0,244 | 688 | 82,735 | 12,798 | 510 | 0,105 | 3555 | 0,533 | 1578 | 76,328 | 15,527 | 1035 | 0,606 | HIGH |
| KRT12.4.L | 8684 | 2,179 | 1784 | 55,232 | 24,793 | 988 | 0,443 | 229187 | 34,381 | 4107 | 38,389 | 22,022 | 1468 | 0,720 | HIGH |
| MAFB.S | 832 | 0,209 | 640 | 83,940 | 12,723 | 507 | 0,057 | 17593 | 2,639 | 2516 | 62,256 | 15,917 | 1061 | 0,714 | HIGH |
| MSX1.L | 12232 | 3,070 | 2398 | 39,824 | 18,168 | 724 | 0,580 | 13925 | 2,089 | 2509 | 62,361 | 13,426 | 895 | 0,665 | HIGH |
| MSX2.L | 4647 | 1,166 | 1638 | 58,896 | 20,627 | 822 | 0,470 | 2651 | 0,398 | 1423 | 78,653 | 14,506 | 967 | 0,369 | HIGH |
| MYL1.L | 1541 | 0,387 | 609 | 84,718 | 8,708 | 347 | 0,518 | 152127 | 22,821 | 2807 | 57,891 | 16,412 | 1094 | 0,932 | HIGH |
| MYOD1.S | 25513 | 6,402 | 2680 | 32,748 | 18,745 | 747 | 0,687 | 30254 | 4,539 | 2149 | 67,762 | 13,921 | 928 | 0,896 | HIGH |
| NOG.L | 3422 | 0,859 | 860 | 78,419 | 12,773 | 509 | 0,726 | 4521 | 0,678 | 1815 | 72,772 | 15,032 | 1002 | 0,492 | HIGH |
| OLIG3.S | 326 | 0,082 | 203 | 94,906 | 4,291 | 171 | 0,310 | 2658 | 0,399 | 1330 | 80,048 | 14,806 | 987 | 0,379 | HIGH |
| OLIG4.L | 910 | 0,228 | 662 | 83,388 | 13,124 | 523 | 0,212 | 2624 | 0,394 | 1650 | 75,248 | 17,762 | 1184 | 0,246 | MEDIUM |
| OTX2.S | 1013 | 0,254 | 738 | 81,481 | 14,630 | 583 | 0,291 | 12913 | 1,937 | 2606 | 60,906 | 15,212 | 1014 | 0,783 | HIGH |
| PAX3.S | 4085 | 1,025 | 1249 | 68,657 | 17,240 | 687 | 0,619 | 5083 | 0,763 | 1576 | 76,358 | 13,501 | 900 | 0,601 | HIGH |
| PAX6.S | 1088 | 0,273 | 804 | 79,824 | 15,533 | 619 | 0,136 | 10252 | 1,538 | 2295 | 65,572 | 18,617 | 1241 | 0,773 | HIGH |
| PAX7.L | 179 | 0,045 | 169 | 95,759 | 3,990 | 159 | 0,030 | 2915 | 0,437 | 1510 | 77,348 | 15,662 | 1044 | 0,346 | HIGH |
| PITX1.L | 739 | 0,185 | 647 | 83,764 | 14,178 | 565 | 0,009 | 42846 | 6,428 | 2339 | 64,911 | 16,937 | 1129 | 0,890 | HIGH |
| RAX.L | 743 | 0,186 | 601 | 84,918 | 12,271 | 489 | 0,069 | 11695 | 1,754 | 1474 | 77,888 | 12,061 | 804 | 0,851 | HIGH |
| SIX1.L | 364 | 0,091 | 313 | 92,146 | 6,851 | 273 | 0,105 | 45096 | 6,765 | 4316 | 35,254 | 13,306 | 887 | 0,748 | HIGH |
| SNAI1.S | 18157 | 4,556 | 2953 | 25,897 | 17,340 | 691 | 0,503 | 53031 | 7,955 | 5051 | 24,227 | 17,372 | 1158 | 0,644 | HIGH |
| SNAI2.L | 985 | 0,247 | 787 | 80,251 | 16,010 | 638 | 0,048 | 13020 | 1,953 | 2019 | 69,712 | 15,122 | 1008 | 0,710 | HIGH |
| SOX10.L | 691 | 0,173 | 583 | 85,370 | 12,296 | 490 | 0,025 | 14858 | 2,229 | 1428 | 78,578 | 11,101 | 740 | 0,818 | HIGH |
| SOX17A.S | 11293 | 2,834 | 1844 | 53,726 | 19,448 | 775 | 0,597 | 17523 | 2,629 | 1895 | 71,572 | 11,446 | 763 | 0,746 | HIGH |
| SOX2.L | 16198 | 4,065 | 2147 | 46,123 | 19,373 | 772 | 0,720 | 30376 | 4,557 | 2948 | 55,776 | 13,531 | 902 | 0,843 | HIGH |
| SOX3.S | 34338 | 8,617 | 2961 | 25,696 | 17,641 | 703 | 0,677 | 61509 | 9,227 | 3283 | 50,750 | 15,257 | 1017 | 0,848 | HIGH |
| SOX8.L | 4058 | 1,018 | 1558 | 60,903 | 20,000 | 797 | 0,401 | 19681 | 2,952 | 2046 | 69,307 | 16,112 | 1074 | 0,784 | HIGH |
| SOX9.S | 3953 | 0,992 | 1369 | 65,646 | 16,763 | 668 | 0,407 | 19746 | 2,962 | 3079 | 53,810 | 17,897 | 1193 | 0,644 | HIGH |
| TBXT.S | 1493 | 0,375 | 1101 | 72,371 | 20,452 | 815 | 0,071 | 955 | 0,143 | 814 | 87,789 | 10,531 | 702 | 0,049 | LOW |
| TFAP2A.L | 6384 | 1,602 | 1880 | 52,823 | 21,305 | 849 | 0,519 | 22710 | 3,407 | 3099 | 53,510 | 15,932 | 1062 | 0,726 | HIGH |
| TFAP2C.L | 20322 | 5,100 | 3029 | 23,990 | 19,674 | 784 | 0,511 | 42804 | 6,421 | 3594 | 46,085 | 17,282 | 1152 | 0,737 | HIGH |
| TP63.L | 877 | 0,220 | 695 | 82,560 | 14,454 | 576 | 0,095 | 11627 | 1,744 | 2088 | 68,677 | 14,611 | 974 | 0,452 | HIGH |
| VENTX2.1.L | 7630 | 1,915 | 2716 | 31,844 | 23,940 | 954 | 0,289 | 2665 | 0,400 | 1717 | 74,242 | 17,717 | 1181 | 0,142 | HIGH |
| ZIC1.S | 10273 | 2,578 | 2039 | 48,833 | 21,380 | 852 | 0,677 | 12273 | 1,841 | 2371 | 64,431 | 17,207 | 1147 | 0,723 | HIGH |

**Zhou et al., Figure S1**

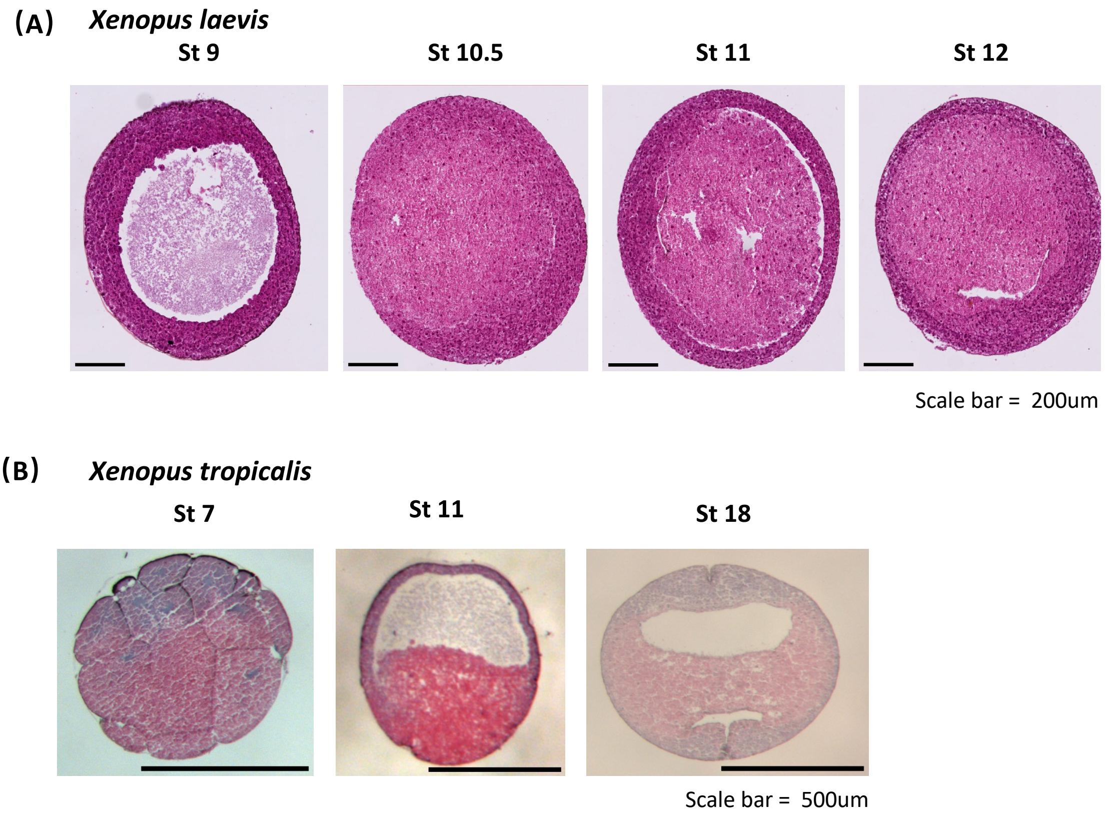

#### Zhou et al., Figure S2

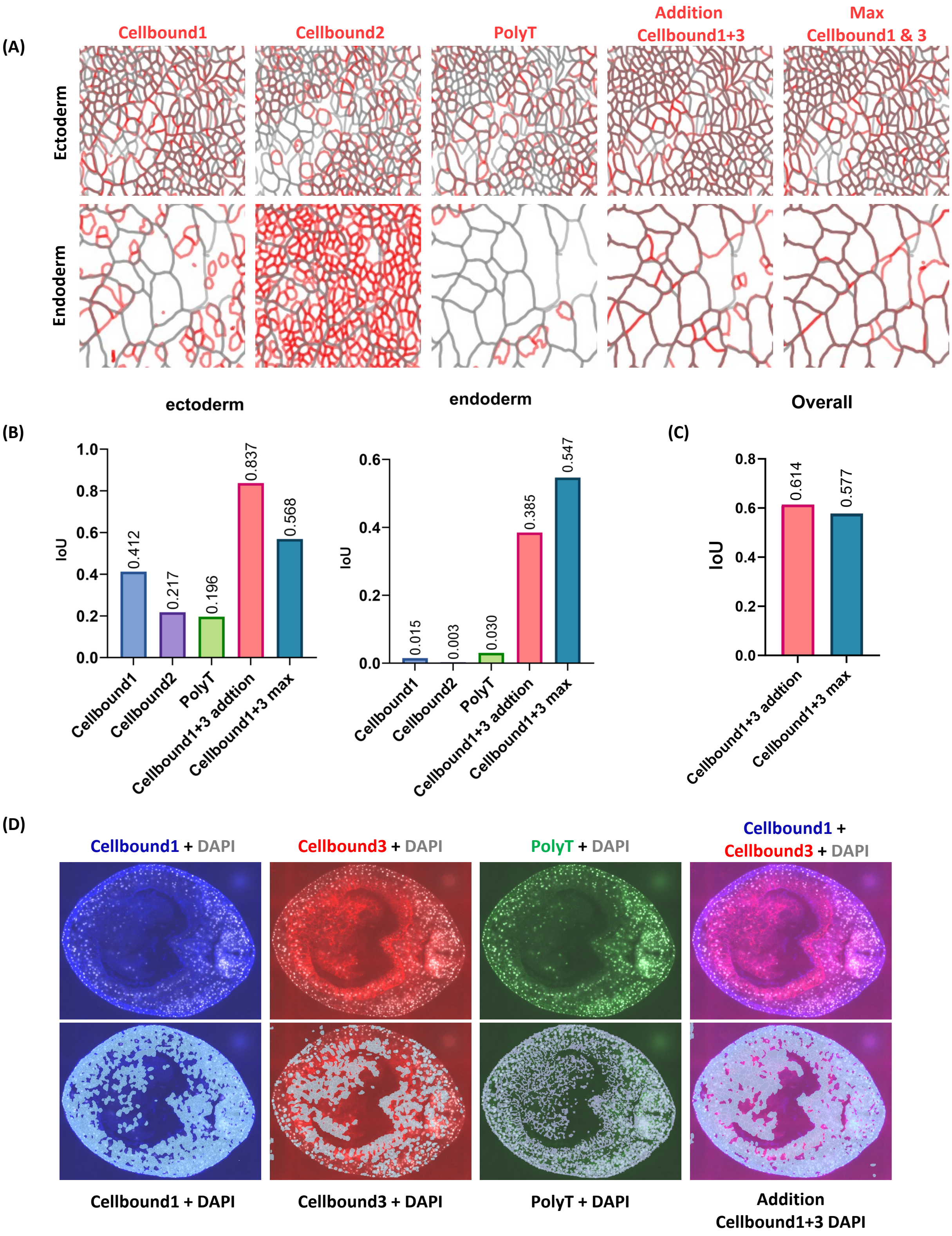

**Zhou et al., Figure S3**

Cell segmentation (blue on the right) for stage 12.5 to stage 22 *Xenopus laevis* embryos.

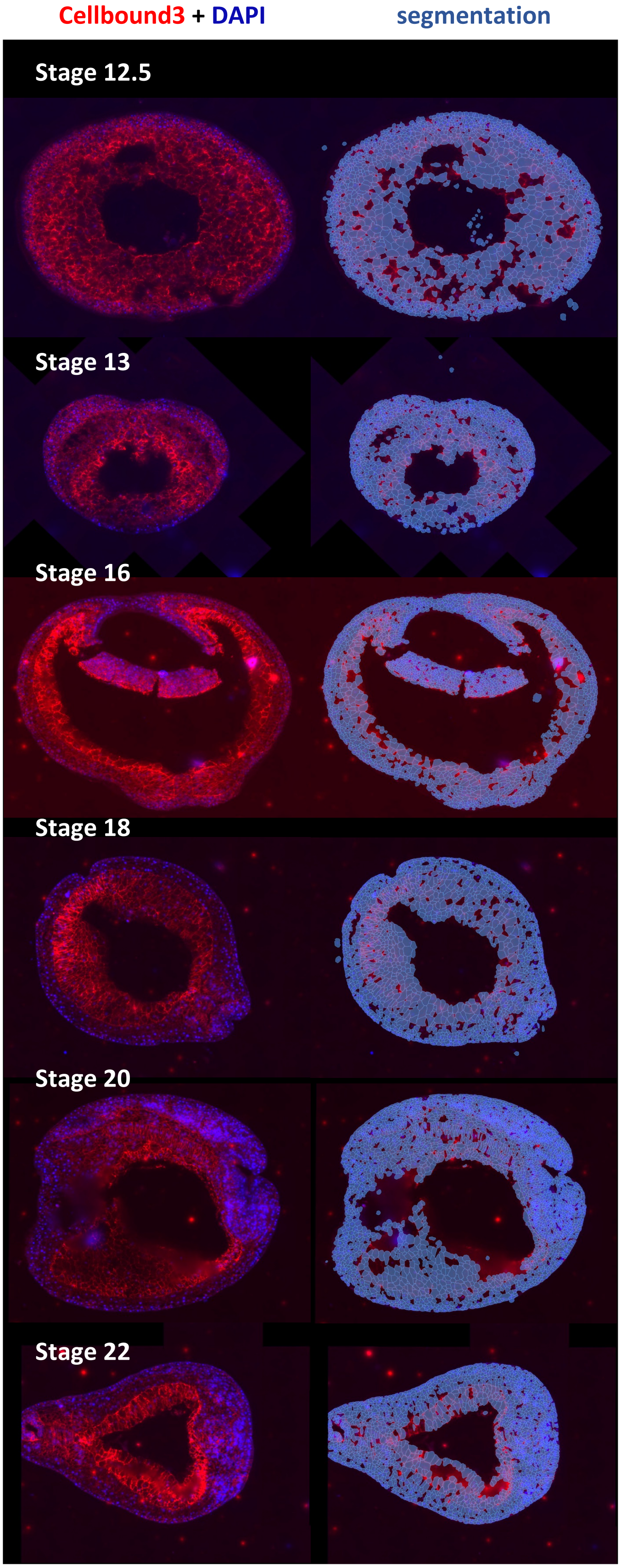

#### Zhou et al., Figure S4

**(A) Transcripts captured inside the cells by Model 1 and Model 2**

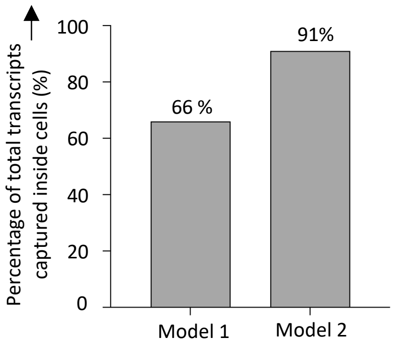


**(B) Comparison of Cell volume distribution by Cell type between Model 1 and Model 2**

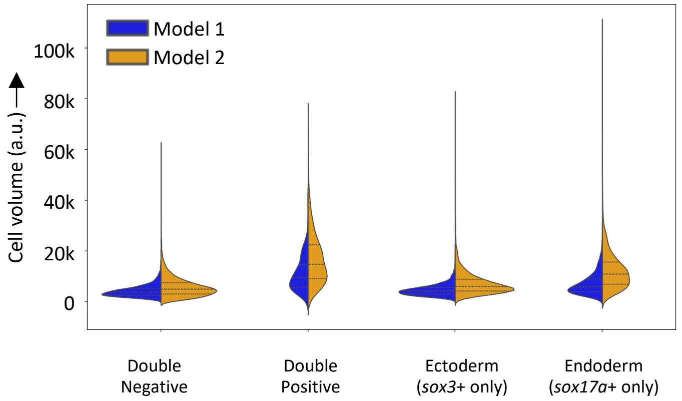


**(C) *sox17*+ Endoderm Cell Distribution by Segmentation Model**

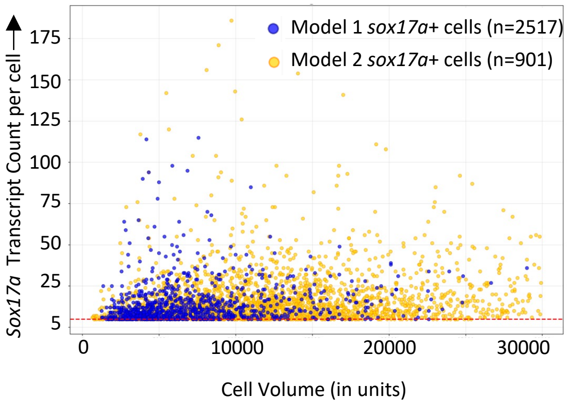

Zhou et al., Figure S5

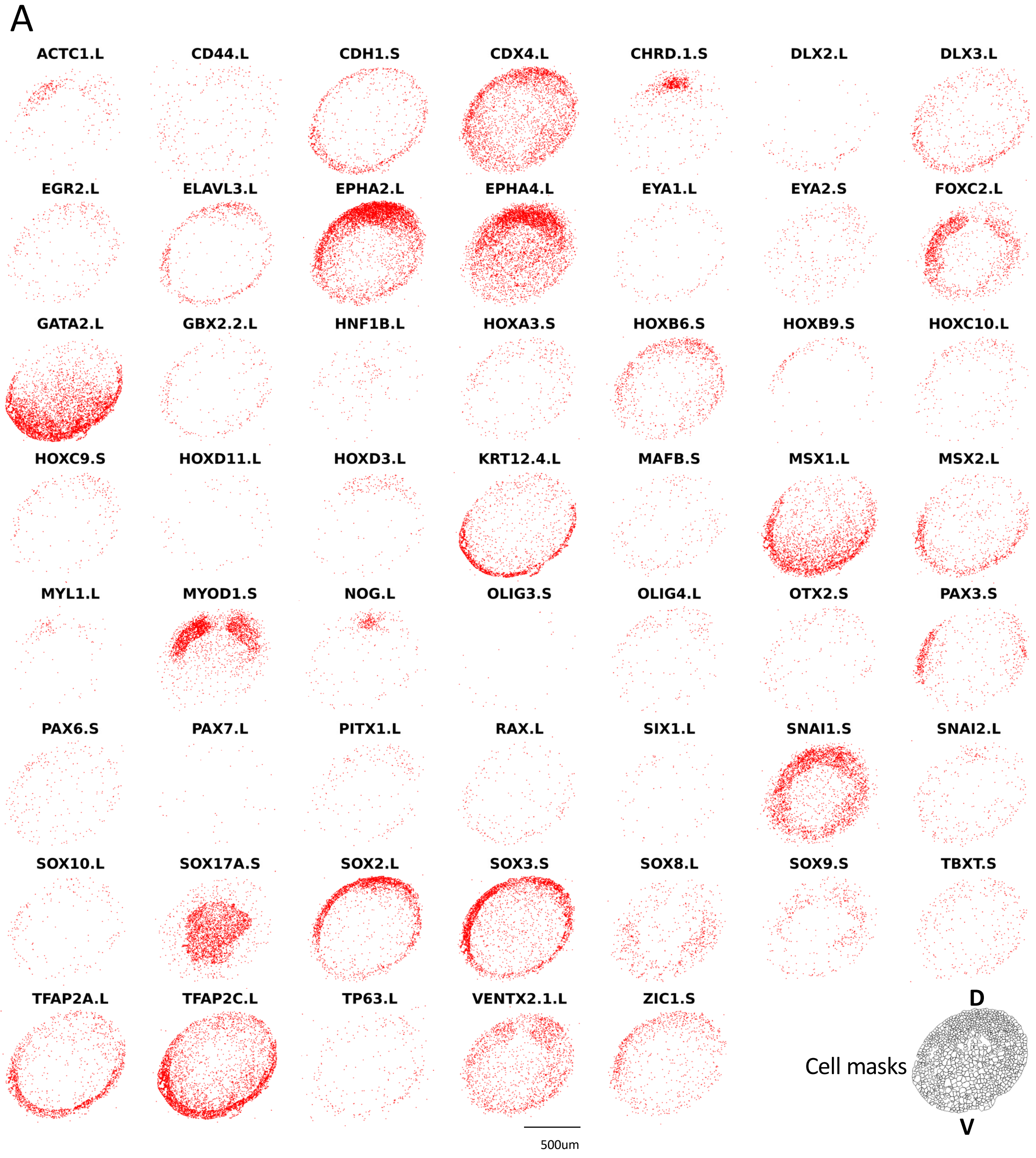

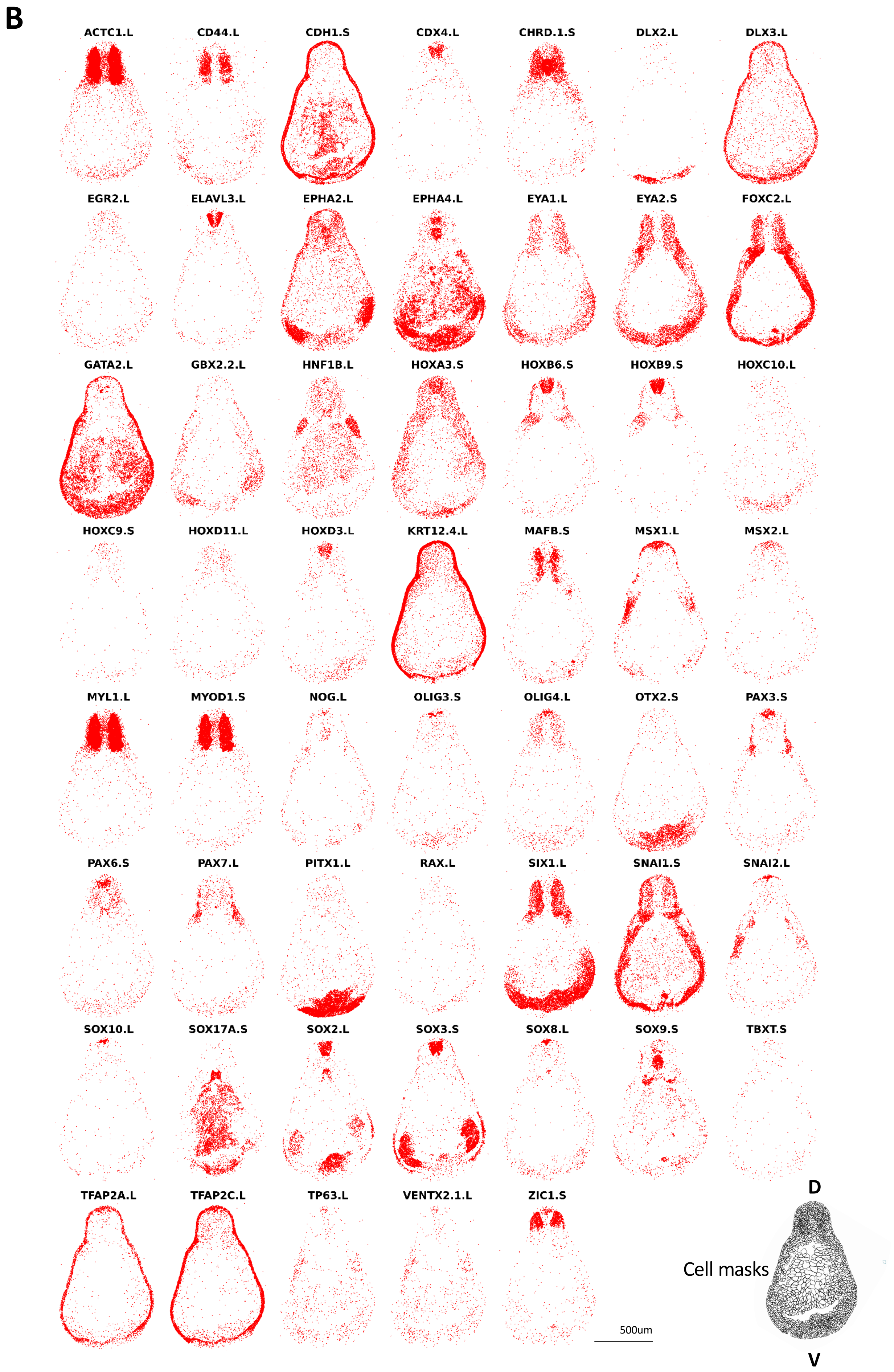
B
ACTC1.L
CD44.L
CDH1.S
CDX4.L
CHRD.1.S
DLX2.L
DLX3.L
EGR2.L
ELAVL3.L
EPHA2.L
EPHA4.L
EYA1.L
EYA2.S
FOXC2.L
GATA2.L
GBX2.2.L
HNF1B.L
HOXA3.S
HOXB6.S
HOXB9.S
HOXC10.L
HOXC9.S
HOXD11.L
HOXD3.L
KRT12.4.L
MAFB.S
MSX1.L
MSX2.L
MYL1.L
MYOD1.S
NOG.L
OLIG3.S
OLIG4.L
OTX2.S
PAX3.S
PAX6.S
PAX7.L
PITX1.L
RAX.L
SIX1.L
SNAI1.S
SNAI2.L
SOX10.L
SOX17A.S
SOX2.L
SOX3.S
SOX8.L
SOX9.S
TBXT.S
TFAP2A.L
TFAP2C.L
TP63.L
VENTX2.1.L
ZIC1.S
500um
D
Cell masks
V

Zhou et al., Figure S6

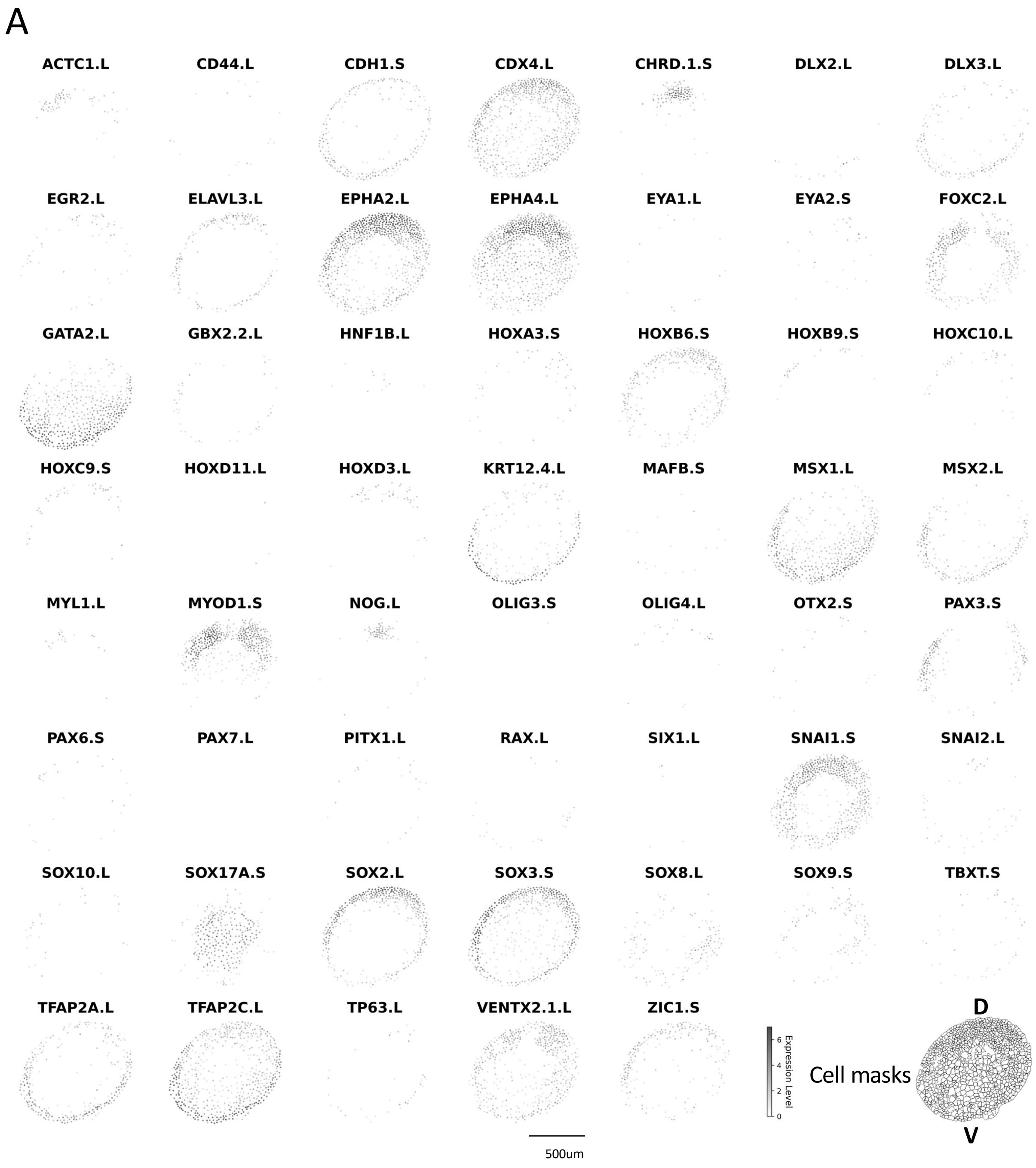

Zhou et al., Figure S6 (continued)

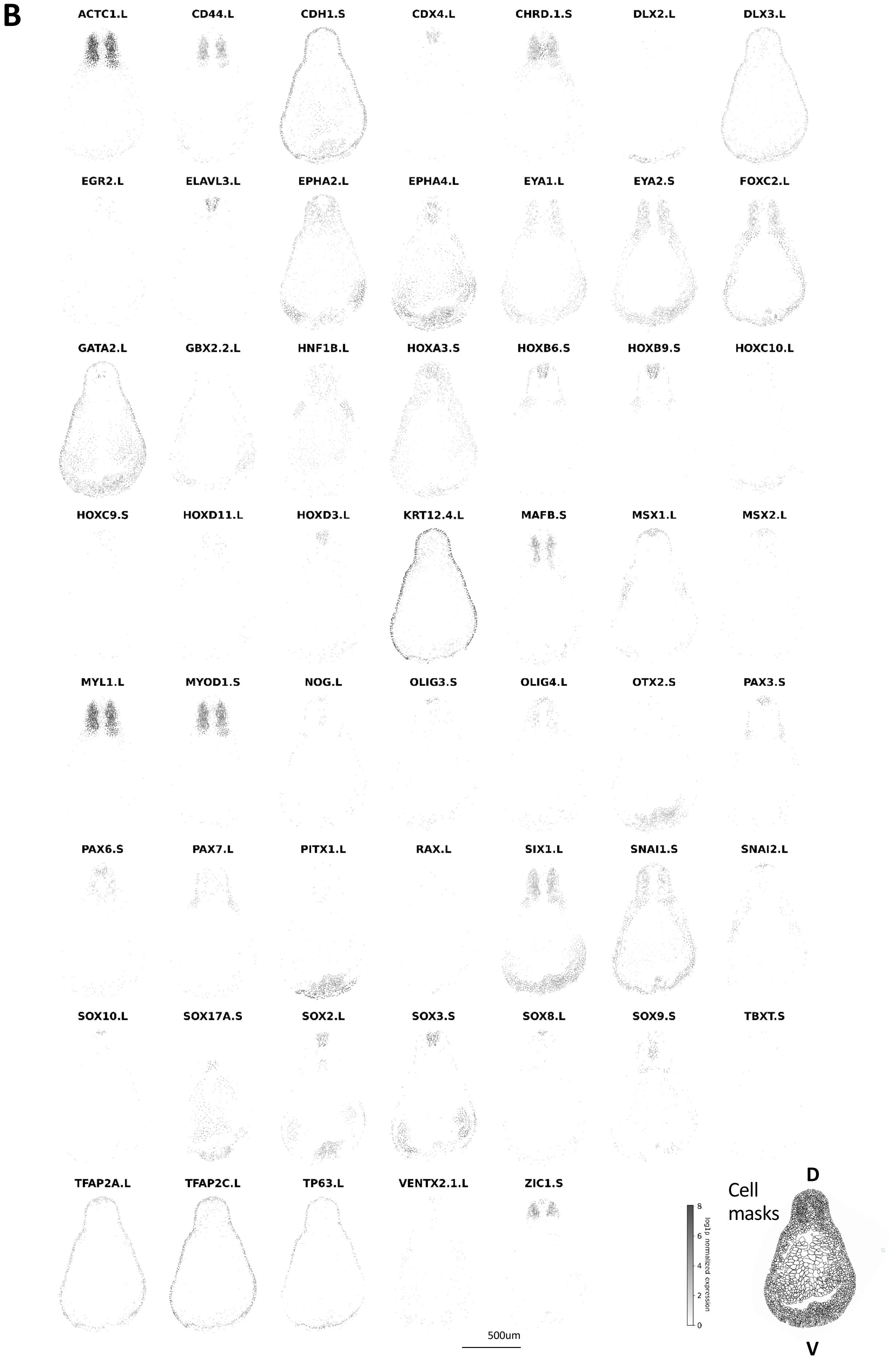

### Zhou et al., Figure S7

#### (A) UMAP plots of gene-based clusters

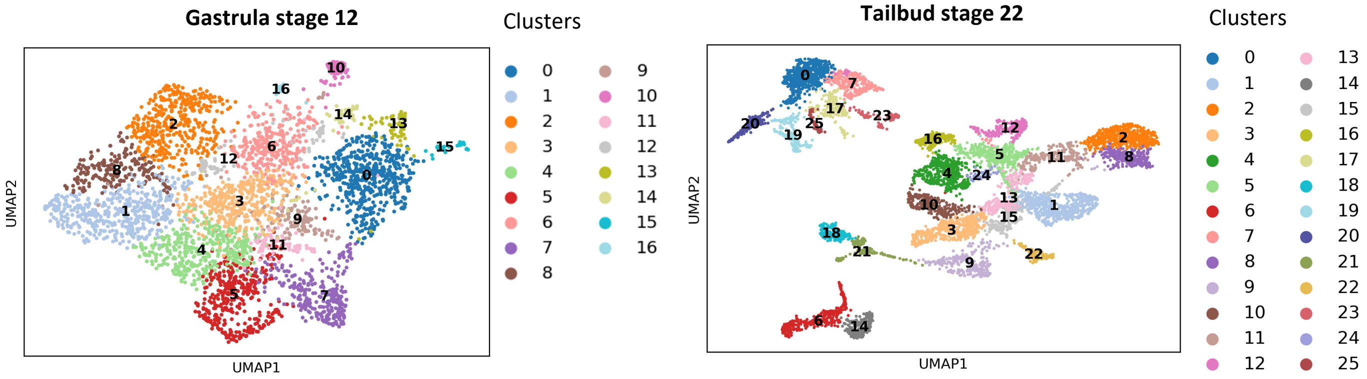


#### (B) Expression of top 3 genes per cluster

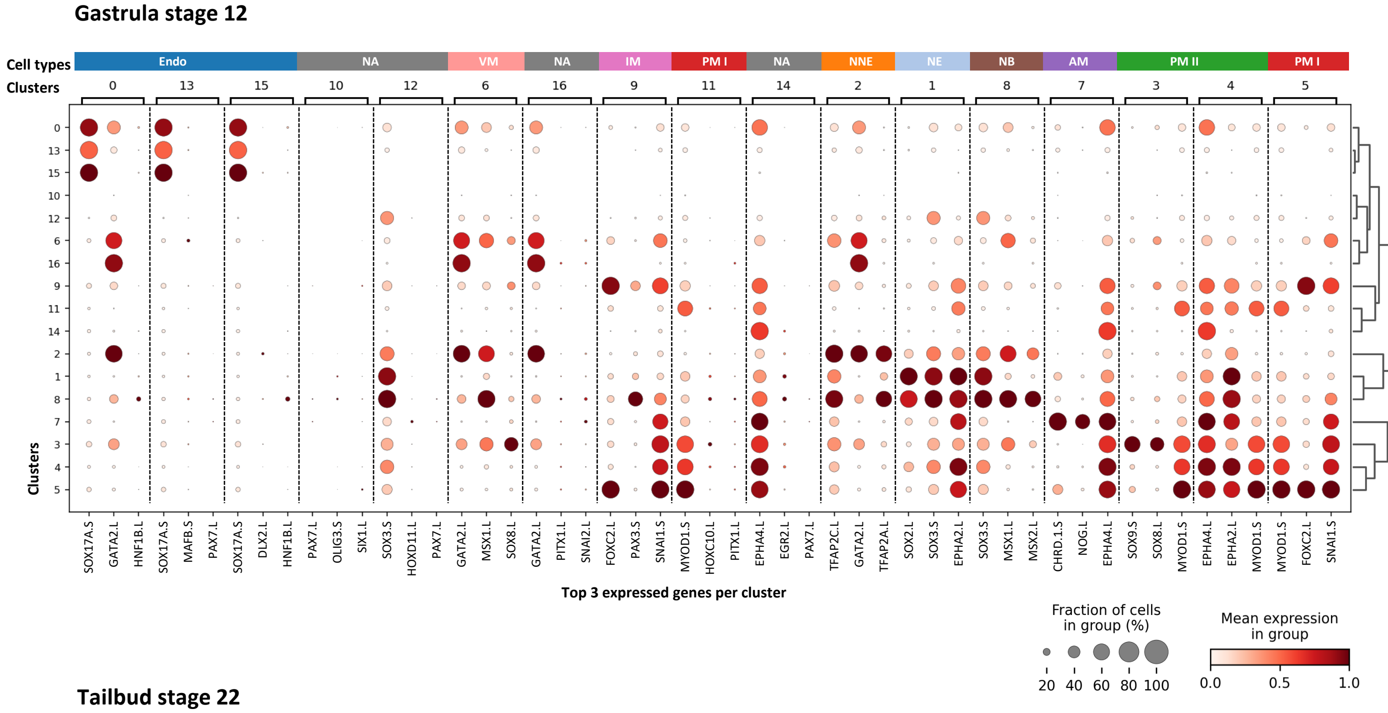


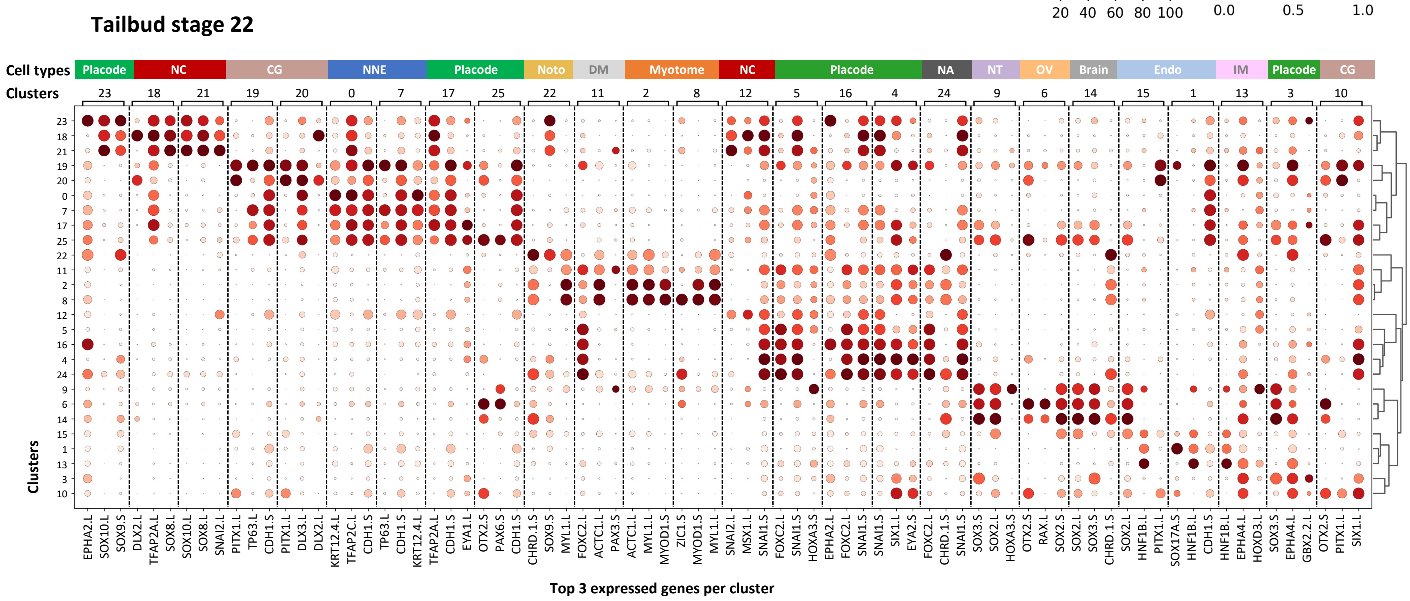

#### Zhou et al., Figure S8

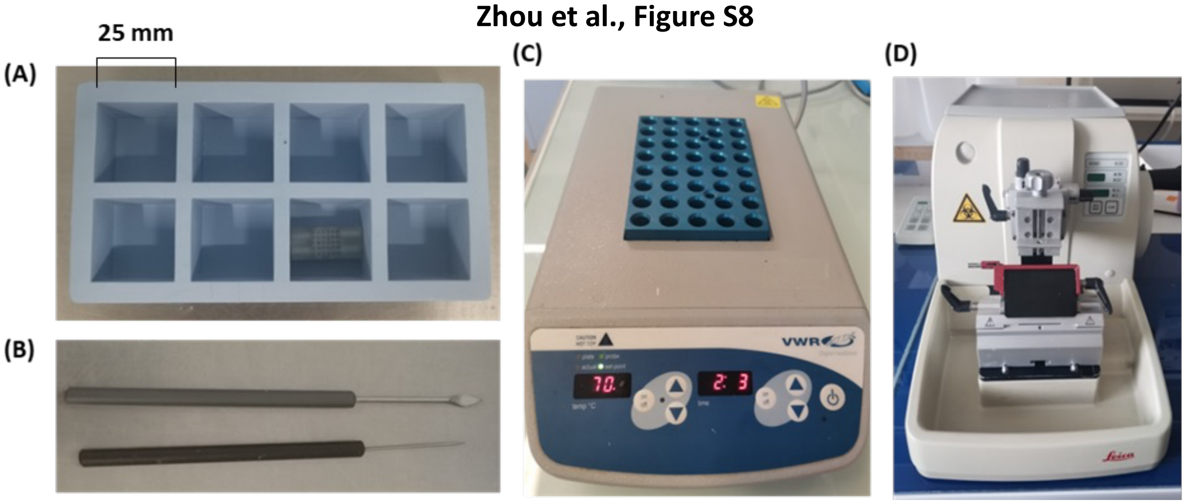


#### Zhou et al., Figure S9

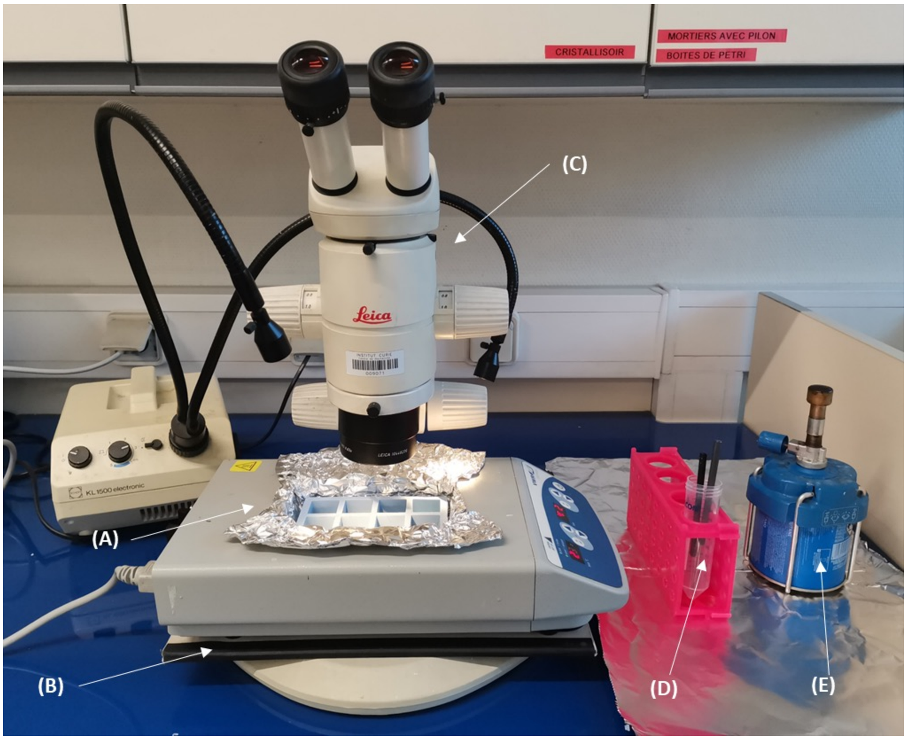


#### Zhou et al., Figure S10

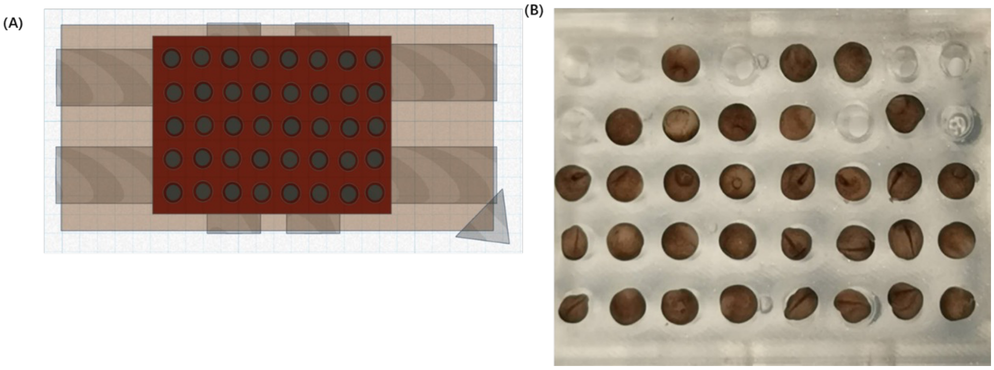
